# Multiomic Analysis of Adult Diapause in *Drosophila melanogaster* Identifies Hallmarks of Cellular Quiescence

**DOI:** 10.1101/2022.12.21.521440

**Authors:** Corinne Hutfilz

**Author notes:** Correspondence: Corinne Hutfilz. **Author Contributions:** C.H. is the sole contributor to this work.

## Abstract

Age is a fundamental aspect of biology that underlies the efficacy of a broad range of functions. Identifying determinants for how quickly or slowly we age will contribute greatly to our understanding of age as a modifier of overall health, particularly to the advancement of therapeutic interventions designed to mitigate or delay age-associated disorders. While much work has been devoted to the study of genetic or pharmacological interventions that extend lifespan, this approach does not necessarily recapitulate the physiological profile of naturally long-lived individuals. Diapause and diapause-like states constitute natural, inducible and evolutionarily conserved examples of lifespan plasticity that are well-suited to serve as physiologically accurate models of longevity. Here, we leveraged a metabolically critical signaling organ in Drosophila, the fat body, to examine diapause-associated transcription in the context of chromatin accessibility and the regulation of lifespan. Through a combination of ATAC-seq and RNA-seq, our observations suggest chromatin is globally reorganized in diapause and may assume a poised conformation to facilitate the rapid transcription of pro-development genes upon diapause termination. We found particular significance of GAF, NELF, and RNA polymerase III in this context. Congruently, transcription during diapause appears to favor many processes supporting the maintenance of cellular quiescence and the inhibition of differentiation. Our data are consistent with a model wherein diapause induces cellular quiescence in the fat body, as was additionally supported through fluorescent microscopy and comparison with public ChIP-seq data for developmentally juvenile files. This work opens the possibility that longevity in diapause may be partially determined through a lack of mitogenic signaling from the quiescent niche, concurrent with changes to the hormonal and immunological profiles that skew metabolism towards tissue maintenance.

## Introduction

Adult insect diapause is a physiological adaptation to unfavorable conditions characterized by delayed development, metabolic adaptation, reproductive dormancy and extreme longevity. Although it is best studied in contexts of development and seasonality, diapause represents a unique model system to understand the mechanisms controlling our rates of aging. Lifespan extends dramatically on induction of diapause. Upon diapause termination, this extension is reversed to pre-diapause lengths without impinging on the organism’s total lifespan at normal conditions. Aging is therefore suspended for the duration of diapause, shielding the species from dying out during seasons that would otherwise prevent the generation of successful progeny. This remarkable plasticity implies a change in the molecular basis for lifespan. Using diapause as a model, these mechanisms can be understood in their evolutionary context to improve our knowledge of the contribution of age to all other aspects of physiology. This has great significance for the advancement of therapies in the course of geriatric care, as well as the prevention of age-associated disease.

Variations of diapause are widely conserved across taxa. Mammals such as bats, squirrels and bears enter hibernation in response to similar environmental conditions that induce insect diapause (Hussein et al., 2020; Wu & Storey, 2021). Ectothermic animals use estivation to limit activity during hot or dry periods, while many rodents and marsupials additionally exhibit an embryonic diapause at the blastocyst stage by delaying implantation into the uterine wall (Staples, 2016; Deng et al., 2018; Ptak et al., 2012). These variations possess distinguishing features both physiologically and in terms of stimulus, but also employ conserved metabolic strategies to prolong life, mitigate tissue damage, and maintain developmental plasticity for the duration that environmental conditions remain unfavorable. The branching nature of diapause-like adaptations suggests a foundational biological program for lifespan plasticity that underlies the diversity of lifespan we observe in nature.

This is perhaps best demonstrated through the success of genetic, pharmaceutical, or dietary interventions that extend lifespan in laboratory settings. Modifications to downstream effectors of insulin/insulin-like signaling (IIS), ribogenesis or autophagy, for example, are broadly characteristic of diapause physiology and have been additionally shown to extend lifespan in non-diapause laboratory settings (Partridge et al., 2011; Tiku & Antebi, 2018; MacInnes, 2016; Bjedov et al., 2010; Nadeau et al., 2022; Lenz et al., 2021; Hofmann & Hand, 1994). Organisms in diapause are longer-lived than non-diapausing organisms even with these modifications, however, suggesting that individual disruptions in the pathways known to regulate lifespan represent partial diapause phenotypes. The lifespan extension observed, under this model, would therefore be attributable to the activation of a conserved ‘diapause response,’ shared across taxa. Supporting this idea, it is often reported that other diapause-like traits, such as increased stress resistance and lowered reproductive propensity, accompany lifespan extension achieved through laboratory means (Partridge et al., 2011).

In *Drosophila*, reproductive maturity is directed by juvenile hormone (JH). The degradation of circulating JH is a typical feature of diapause that causes developmental regression of the ovaries. Loss of JH is also associated with other diapause-like traits, such as increased expression of immune genes, longevity, slow growth and the downregulation of IIS (Hutfilz, 2022). Considering these effects, it is reasonable to consider that diapause physiology, or the extension of lifespan in diapause more specifically, may be reducible to the loss of JH. However, little is known about the molecular functions of JH in diapause, and past work has shown that the loss of JH signaling is not sufficient to fully recapitulate diapause physiology or lifespan (Jindra et al., 2013). Organisms in diapause are much longer lived than organisms that have been made unable to synthesize JH through genetic or surgical means. Transcription-based studies (qPCR, RNA-seq) have additionally shown that IIS profiles are dissimilar between organisms in diapause and those unable to synthesize JH (Yamamoto et al., 2013; Kučerová et al., 2016). Furthermore, some instances of diapause entail periods of fluctuation in JH levels. Natural increases in JH over time do not terminate diapause and the organism remains extremely long-lived. For these reasons, the loss of JH can be considered a contributing, yet confounding factor when studying mechanisms of lifespan plasticity in diapause. It is therefore important to understand the role of JH both in isolation and in context of the greater longevity program.

A separate motivation for this design is the fact that JHs, while common in insects, are absent from other taxonomic classes (Goodman & Granger, 2005). Studying diapause-associated mechanisms of lifespan plasticity that function outside the contribution of JH will therefore not only disentangle JH-dependent effects, but will afford much greater clarity to the ways in which conclusions from diapause research translate to species without JH. The conservation of diapause/diapause-like states across taxa suggests that mechanisms controlling diapause-associated lifespan might be applicable to mammals. This provides motivation to study modes of longevity represented through partial diapause phenotypes (mutations in IIS or JH loss, for example), but perhaps more importantly, to study diapause *qua* diapause, with an eye towards pathways orthologous to humans.

Previous transcriptomic work has described the molecular processes underlying diapause physiology in broad terms. However, these experiments have most often been performed using whole body preparation, which complicates interpretation and risks the underrepresentation or overrepresentation of key processes carried out by specific tissues. Additionally, there has been very little work done to examine upstream regulatory mechanisms responsible for mediating the diapause transcriptome and its outcomes for longevity. In this study, we therefore chose to investigate the chromatin architecture associated with a critical signaling tissue in diapause, the fat body, because the combination of these two approaches represents a convergence of many upstream factors controlling transcription.

The fat body is a signaling organ that coordinates systemic change to physiology through the integration of nutritional and endocrine signals from other tissues. The fat body regulates diverse processes such as growth, longevity, energy storage and the innate immune response, among other functions contributing to metabolic homeostasis. Many of these functions are regulated through the fat body’s control of insulin/insulin-like signaling (IIS). The fat body both receives and secretes different insulin-like peptides (ILPs) to modulate growth according to environmental conditions, making it a central tissue in diapause entrance, maintenance and termination.

In holometabolous insects such as *Drosophila*, the development of the fat body progresses through two phases. The first group of cells comprising the fat body, known as the larval fat, is derived from the mesoderm and provides necessary energy and amino acids for metamorphosis during the non-feeding pupal stage of development. The larval fat also performs signaling functions to initiate metamorphosis and sexual maturation. Sensors of adiposity such as Upd2 and apolipophorin control the larva’s ability to link nutrition and development through the stimulation of ribosome biogenesis in the prothoracic gland (Juarez-Carreño et al., 2021). Larval fat body cells persist from the larval stages into early adulthood, whereupon they undergo apoptosis within the first two to four days after eclosion from the pupal case (Rizki, 1978). The larval fat body is simultaneously replaced by adult fat body cells, whose tissue of origin remains unknown, but are possibly derived from ectodermal cells of the dorsal thoracic imaginal discs, or by histoblasts associated with the larval body wall (Hoshizaki, 2005; Hoshizaki et al., 1994). Histoblasts are specified during embryonic stages of development and remain in a state of G2 cell cycle arrest during subsequent larval stages (Verma & Cohen, 2015). Unlike larval tissues that grow primarily via endoreplication, the progenitor cells of adult-specific tissues divide through mitosis (Zielke et al., 2013; Hoshizaki, 2005). Thus, the final stages of development in *Drosophila* depend on mitogenic signals to cue the end of progenitor cell quiescence.

Certain features are shared among instances of cellular quiescence, while others appear to be dependent on species or cell type. Here we will briefly examine these hallmarks. Cell cycle inhibition in quiescence is characterized by distinct processes from other nondividing populations. All quiescent cells must maintain a state of replicative dormancy without sacrificing proliferative potential. This entails the assumption of a “primed” state where the cell remains sensitive to cues that signal cell cycle reentrance. Exit from the cell cycle is mediated through the inhibition of cyclins and cyclin-dependent kinases (CDKs), as well as the E2F1 transcription factor. *Drosophila* transcribe two E2F proteins; E2F2 is primarily repressive and opposes the cell cycle-promoting activity of E2F1 by recruiting proteins of the retinoblastoma (Rb) family to E2F1 target genes (Weng et al., 2003; Øvrebø et al., 2022). In *Drosophila* neuroblasts, Hox gene expression also plays an important role in the induction of quiescence (Tsuji et al., 2008).

The maintenance of the quiescent state is mediated through multiple pathways, such as the absence of soluble mitogenic signals at the quiescent niche and the retention of cell-cell adhesion through cadherins (Gao et al., 2013). Quiescent cells exhibit a downregulation of RNA synthesis, which involves a general decline in the transcription of mRNAs partially dependent on the pausing of RNA polymerase II (RNAPII), as well as the rRNAs, snoRNAs and tRNAs necessary for protein synthesis (Core & Adelman, 2019; Wang & Amoyel, 2022; Gala et al., 2022; van Velthoven & Rando, 2019; Freter et al., 2010; Loayza-Puch et al., 2013). This reduces the cell’s metabolic demand for nucleotides and amino acids. However, quiescence also entails upregulation of genes responsible for preventing differentiation, maintaining cellular homeostasis, and the transduction of mitogenic signals, such that the cell remains primed for rapid return to the cell cycle. Positive regulation of the Notch (N) and Hippo (hpo) signaling pathways is crucial to maintaining undifferentiated cell identity in quiescence (van Velthoven & Rando, 2019; Sood et al., 2022; Bjornson et al., 2012; Ding et al., 2016). The prevention of damage accumulation is another necessary function in quiescence insofar as the cell must preserve its genomic integrity for future replication. This is partially mediated through upregulation of the transcription factor FoxO, which additionally mediates many other aspects of quiescent cell physiology, including induction. FoxO has also been shown to slow aging (Kops et al., 2002; Zhang et al., 2020; Artoni et al., 2017; Demontis & Perrimon et al., 2010).

FoxO opposes the activity of factors that promote growth and cell cycling, such as the mechanistic target of rapamycin (mTOR) (Puig et al., 2003). mTOR is a pro-aging factor that directs proliferation through the sensing of amino acids and the consequent phosphorylation of eukaryotic translation initiation factor 4E (eIF4E)-binding proteins (4E-BP) (Fernandes & Demetriades, 2021; Jewell et al., 2013; Ben-Sahra & Manning, 2017). Transcription of 4E-BP is driven by FoxO. When phosphorylated, 4E-BP releases from mRNA-bound eIF4E, thus enabling translation. It follows that reactivation of quiescent cells can be initiated through reintroduction of amino acids, and by extension, the resumption of protein synthesis (Britton & Edgar, 1998). To counteract this, FoxO additionally regulates the production of enzymes mediating amino acid catabolism (Coller, 2021).

Apart from amino acids, quiescent cell metabolism is typically defined through the downregulation of oxidative phosphorylation in exchange for energy derived from glycolysis and fatty acid oxidation (Marescal & Cheeseman, 2020). Diverse examples from yeast, humans, mice and *Drosophila* show the entry of cellular quiescence is characterized by a preparatory upregulation of fatty acid biosynthesis (Peselj et al., 2022). Catabolism of stored lipids via fatty acid β-oxidation (FAO) facilitates the energy-intensive process of cell cycle reentry in yeast, but appears to also sustain the tricarboxylic acid cycle (TCA) for redox homeostasis during quiescence in mice and human stem cells (Kalucka et al., 2018; Knobloch et al., 2017; Ramosaj et al., 2021). In *Drosophila*, the upregulation of FAO genes was shown to extend lifespan in a FoxO-dependent manner (Lee et al., 2012). Recent evidence suggests that the TCA metabolites derived from FAO are directed towards the synthesis of NADPH, rather than nucleotides, through the action of Notch (Kalucka et al., 2018). This may constitute one mechanism by which Notch maintains cellular quiescence; indeed, low levels of intracellular ROS are another requirement for the quiescent state (Coller, 2019).

However, metabolism is inconsistent in some respects across quiescent cell populations. For example, *Drosophila* hemocyte precursor cells initiate differentiation with upregulated FAO, while the abrogation of FAO induces G2-M arrest. Stem cells from humans and rats can also enter G1 cell cycle exit upon inhibition of fatty acid synthesis, effectively omitting the buildup of lipid droplets that often accompanies quiescence induction (Cornell et al., 1977). Inhibition of fatty acid synthesis also appears to induce arrest in mouse stem and progenitor cells, but prolonged inhibition results in apoptosis, suggesting a continued role for fatty acid synthesis in the maintenance of some quiescent cells (Knobloch et al., 2013). These instances demonstrate that, while certain aspects of quiescent cell metabolism are well-conserved, others are tuned to the specific needs of cell type, cell phase, quiescence duration, and species.

## Results

### Diapause is a model for lifespan plasticity

Lifespan extension is a defining feature of diapause. To validate our ability to induce diapause, we measured the diapause-associated lifespan extension of newly eclosed w^1118^ females by comparing survivorship between standard conditions (25°C, 12:12 light/dark, 40% relative humidity) and diapause-inducing conditions (11°C, 10:14 light/dark, 40% relative humidity). We observed strong diapause-associated lifespan extension over the course of 35 days, at which point we removed flies from diapause-inducing conditions and placed them at standard conditions to test whether flies would resume normal lifespan without detriment, another key feature of diapause (fig. 1A). Indeed, upon removal from diapause-inducing conditions, flies aged as though the time spent in diapause was inconsequential (fig. 1A subpanel). The total lifespan extension observed was highly significant (p < 2.2e-16, D = 0.8705, two-sample Kolmogorov-Smirnov test).

**Figure 1:**
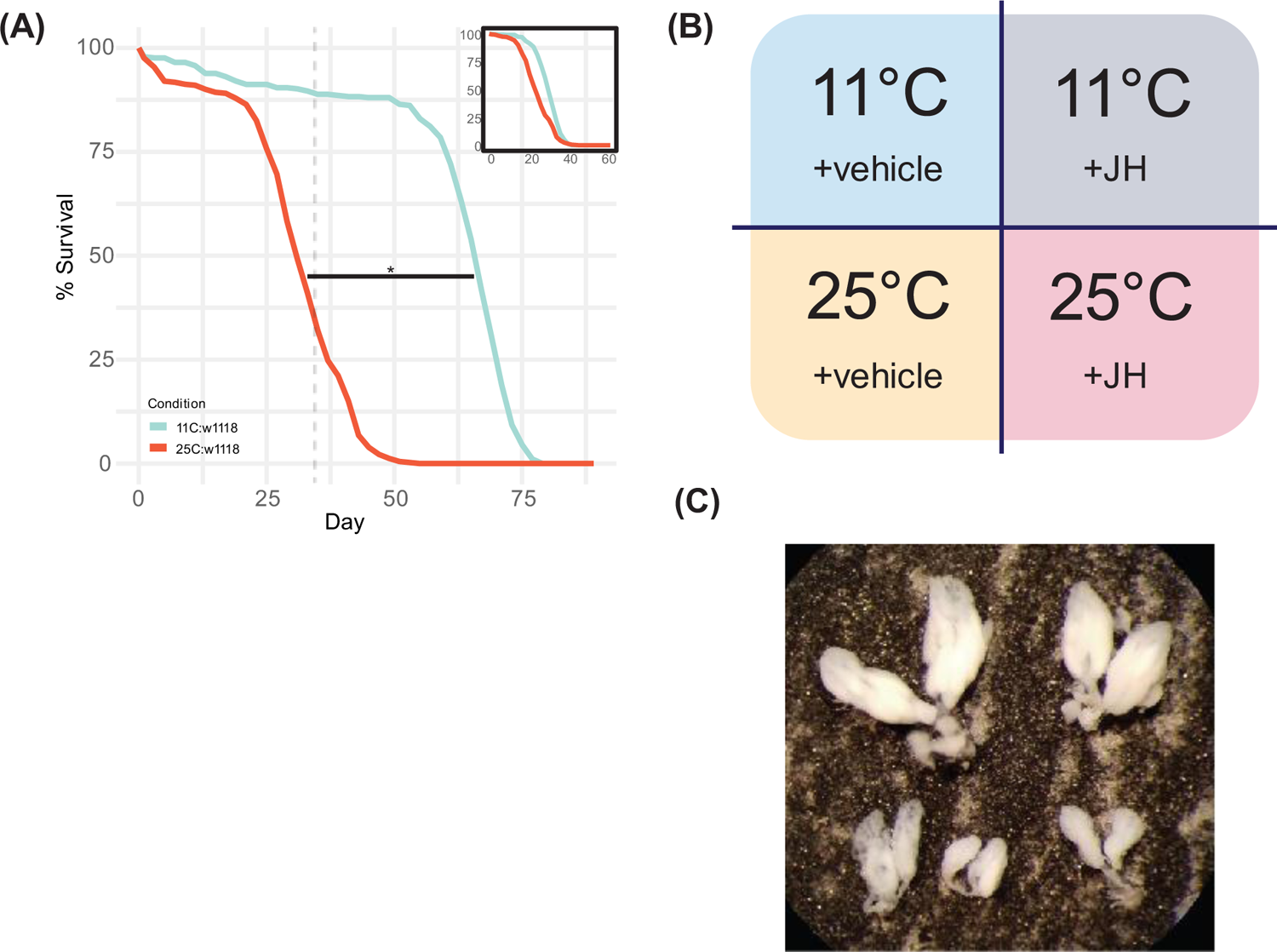
Diapause is a model for lifespan plasticity. **(A)** Survivorship of flies kept in standard conditions (red, n = 412) versus diapause-inducing conditions (blue, n = 376). On day 35 (dashed line), flies in diapause-inducing conditions were placed into standard conditions. Lifespan of flies previously in diapause then reverted to pre-diapause length, as indicated by the top-right subpanel, which censors diapause lifespan by removing data from before day 35. Significance determined by two-sample Kolmogorov-Smirnov test: D = 0.8705, p-value < 2.2e-16. **(B)** Experimental design used for ATAC-seq. S-methoprene (JH) supplementation was used to control for effects due to the natural loss of JH biosynthesis in diapause. JH was diluted in 99% ethanol (vehicle). **(C)** Ovaries dissected from w^1118^ flies under standard conditions (with or without supplemental S-methoprene), or diapause-inducing conditions with supplemental S-methoprene, showed reproductively mature morphology (top row), while ovaries from w^1118^ flies in diapause-inducing conditions with ethanol (vehicle) showed developmentally regressed morphology (bottom row).

To begin probing the mechanisms responsible for this plasticity, we performed Assay for Transposase-Accessible Chromatin by sequencing (ATAC-seq: Buenrostro et al., 2015; Corces et al., 2017) on fat body tissue dissected from age-matched female flies living in the following four conditions: standard conditions with 10 μL ethanol (99%) added every 2 days, standard conditions with 10 μL S-methoprene (39.4 mm, AK Scientific, effective dose 0.1 μg) added every 2 days, diapause-inducing conditions with 10 μL ethanol every 2 days, and diapause-inducing conditions supplemented with 10 μL S-methoprene every 2 days (fig. 1B). S-methoprene is a JH analogue and was diluted in 99% ethanol to allow for volatilization as the mode of delivery. These four conditions were chosen to enable the isolation and elimination of effects dependent on the natural loss of JH synthesis that occurs during diapause. The diapause status of each fly was confirmed before tissue collection through observation of ovarian morphology. Flies kept in standard conditions with or without S-methoprene demonstrated the mature, reproductively viable phenotype, along with flies kept in diapause-inducing conditions supplemented with S-methoprene (fig. 1C, top row). Meanwhile, flies kept in diapause-inducing conditions with ethanol demonstrated the immature, reproductively inviable phenotype (fig. 1C, bottom row). To ensure that S-methoprene treatment was properly represented during analysis, only flies with the anticipated ovarian morphology per condition were used in experiments. Notably, flies without anticipated morphology were very rare. No flies with mature ovarian morphology were observed in diapause-inducing conditions supplemented with ethanol. Similarly, very few flies were observed with immature ovarian morphology in standard conditions of either supplement (approximately 1:40), or in diapause-inducing conditions supplemented with S-methoprene (approximately 1:20). Those flies with immature ovarian morphology in standard conditions generally presented other indications of developmental deformity, such as segmentation defects or melanoma, two complications that are common to inbred strains.

ATAC-seq libraries were generated from each of the four conditions and sequenced to an average depth of approximately 26 million reads per sample. Raw reads were then processed to eliminate mitochondrial reads, optical duplicates and blacklisted peaks. Initial quality assessment showed replicates were highly correlated within conditions (minimum r = 0.91 for standard conditions with EtOH, 0.83 for standard conditions with S-methoprene, 0.95 for diapause-inducing conditions with EtOH, and 0.95 for diapause-inducing conditions with S-methoprene, Pearson correlation coefficient), demonstrating the sample preparation was robust (supplemental fig. S.1, supplemental table S1). A Pearson correlation coefficient of r = 0.9 was used as the threshold for selection of replicates for further processing. Only standard conditions with S-methoprene contained replicates below threshold, resulting in two replicates for this condition. Other conditions retained all four replicates. Differentially accessible genomic loci were then derived using DiffBind at a significance threshold of FDR<0.01. Loci were considered to become “accessible in diapause” and JH-independent if reads were significantly enriched in both diapause-inducing conditions (11°C with or without S-methoprene), but not in either of the standard conditions (25°C with or without S-methoprene). Similarly, loci were considered to become “inaccessible in diapause” and JH-independent if reads were significantly enriched in standard conditions but not in diapause-inducing conditions (fig. 2A). The topmost significantly accessible and inaccessible regions are shown here to demonstrate the experimental design through example (fig. 2B, left and right, respectively). The complete list of differentially accessible loci is available (supplemental table S2). Using this classification, 1,205 regions were identified as differentially accessible in diapause. Of these, 513 regions became accessible and 692 became inaccessible, demonstrating an approximately even distribution between chromatin opening and closing (fig. 2C). There was no obvious localization of changes in accessibility at the level of chromosome; regions of differential accessibility were discovered across all chromosomes (fig. 2D), indicating that diapause imposes global chromatin reorganization.

**Figure 2:**
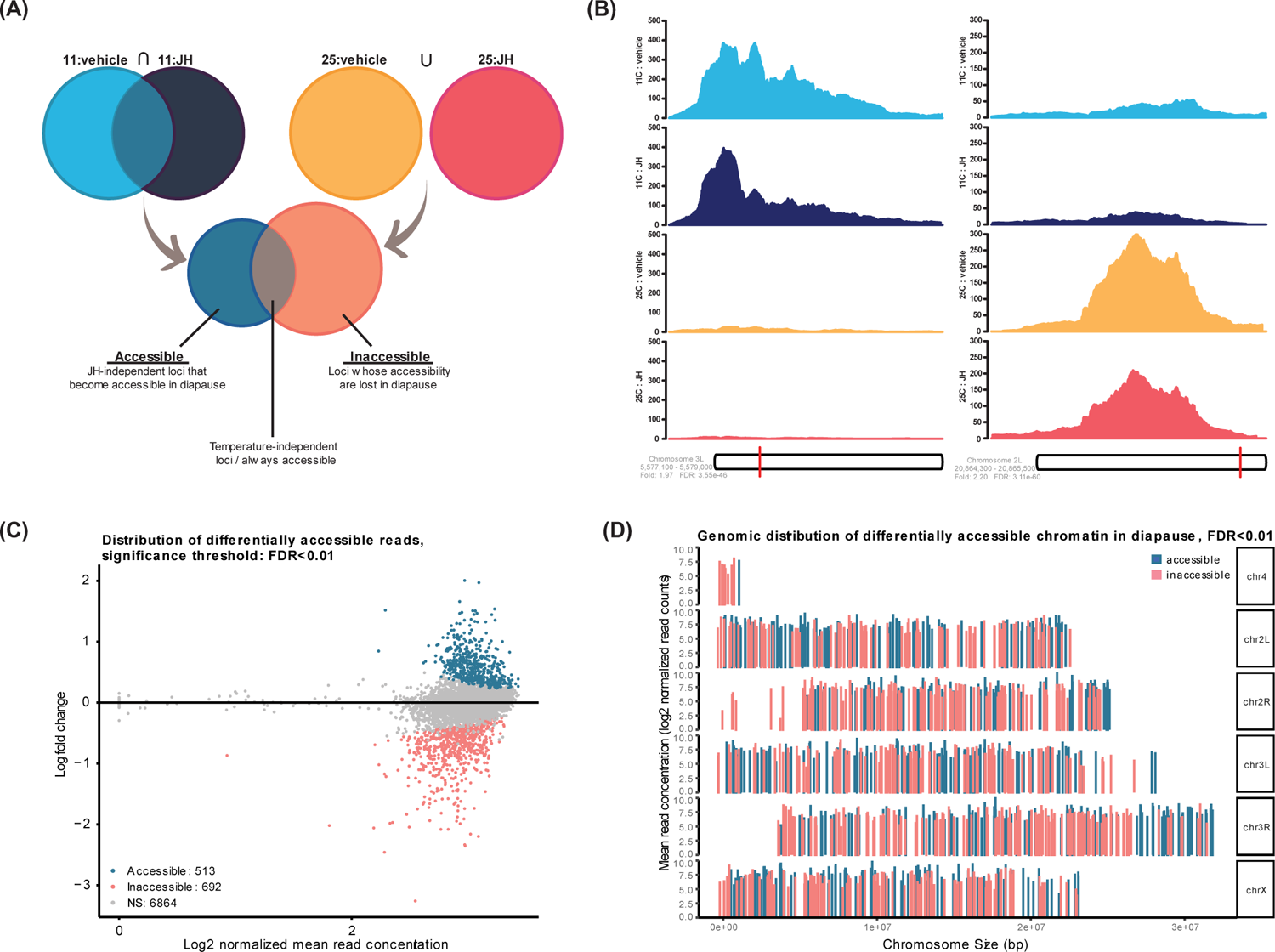
Chromatin is widely reorganized in the diapause fat body independent of JH. **(A)** Filtering method used to derive differentially accessible loci that are independent from the effects of the natural loss of JH biosynthesis in diapause. **(B)** Relative pileup of the most significantly differentially accessible locus that becomes accessible in diapause (left) and inaccessible in diapause (right). **(C)** MA plot showing the distribution of log fold changes for all differentially accessible loci at significance threshold of FDR<0.01. **(D)** Genomic coverage and mean read concentration for differentially accessible loci across chromosomes.

Differentially accessible peaks were then categorized by their alignment to genomic features. Both accessible and inaccessible regions were highly enriched for promoters (59.65% of all accessible regions, 50.29% of all inaccessible regions), as determined by ChIPseeker (Yu et al., 2015) using a ±1 kb window of detection around the transcription start sites of genes to define promoter regions (fig. 3A, B). We also note that euchromatin is enriched at promoters in our data even prior to differential analysis (avg. 75.85%±1.68% standard deviation, supplemental fig. S.2). In light of this information, our analysis illustrates a very large proportion of divergence between the promoter accessibility landscapes of diapause and normal aging. The largest changes in accessibility for non-promoter features post-analysis occur in intronic regions. We did not test whether these intronic regions correspond with enhancer activity.

**Figure 3:**
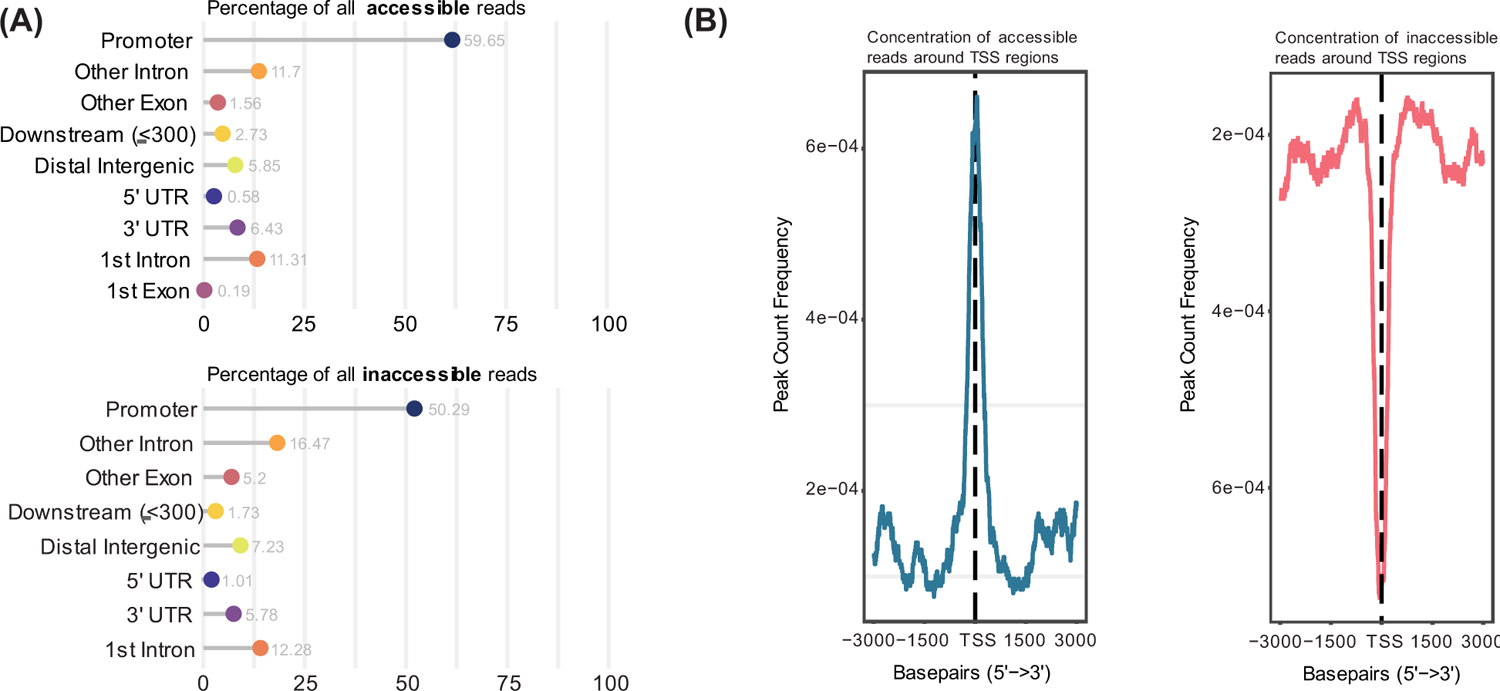
Chromatin reorganization in diapause is enriched at promoter regions. **(A)** Distribution of differentially accessible loci according to the genomic features they map to. Promoters were defined within ±1 kb of a transcription start site. **(B)** Quantification of significantly accessible (left) and inaccessible (right) peak counts within ±3 kb of all transcription start sites.

Together, we show that diapause is regulated by widespread epigenetic changes in the fat body, mainly through the differential accessibility of promoters. We found these changes were distributed evenly throughout the genome, indicating that diapause is accomplished through global chromatin reorganization, as opposed to changes localized to certain chromosomes, or a global skew towards euchromatinization or heterochromatinization.

### Chromatin modifications in diapause facilitate pathways associated with transcriptional poising

We next sought to determine the functional contribution of these differentially accessible regions for diapause physiology and lifespan. We conducted motif analysis using i-cisTarget (Herrmann et al., 2012) under default parameters. To ensure that results were robust, motif analysis was also performed using HOMER (default parameters, fragment size = 200; Heinz et al., 2010) for comparison (supplemental figs. S.3, S.4, supplemental tables S3, S4). Top results were similar between the two platforms, though results from i-cisTarget generally presented more species specificity for *Drosophila* and so are presented here for this reason.

Our analysis showed highly enriched accessibility for the GAGAG motif known in *Drosophila* to bind a suite of factors required for developmental patterning and morphogenesis (fig. 4A). GAGAG motifs are abundantly found in the promoters of genes whose expression is stalled in anticipation of internal or environmental cues that catalyze the reactivation of transcription and consequent progression of development (Qi et al., 2022; Slattery et al., 2014). This stalling occurs via Negative Elongation Factor (NELF), which localizes to RNAPII-loaded promoters via recruitment from GAF (Li et al., 2013; Li & Gilmour et al., 2013; Lee et al., 2008). GAF is well-known to bind the GAGAG motif (van Steensel et al., 2003). The motif is also found outside promoter regions in Polycomb response elements, enhancers, and within the boundary regions of topological domains (Greenberg & Schedl, 2001). The binding motif for the Polycomb catalytic component, E(z), was also significantly accessible.

**Figure 4:**
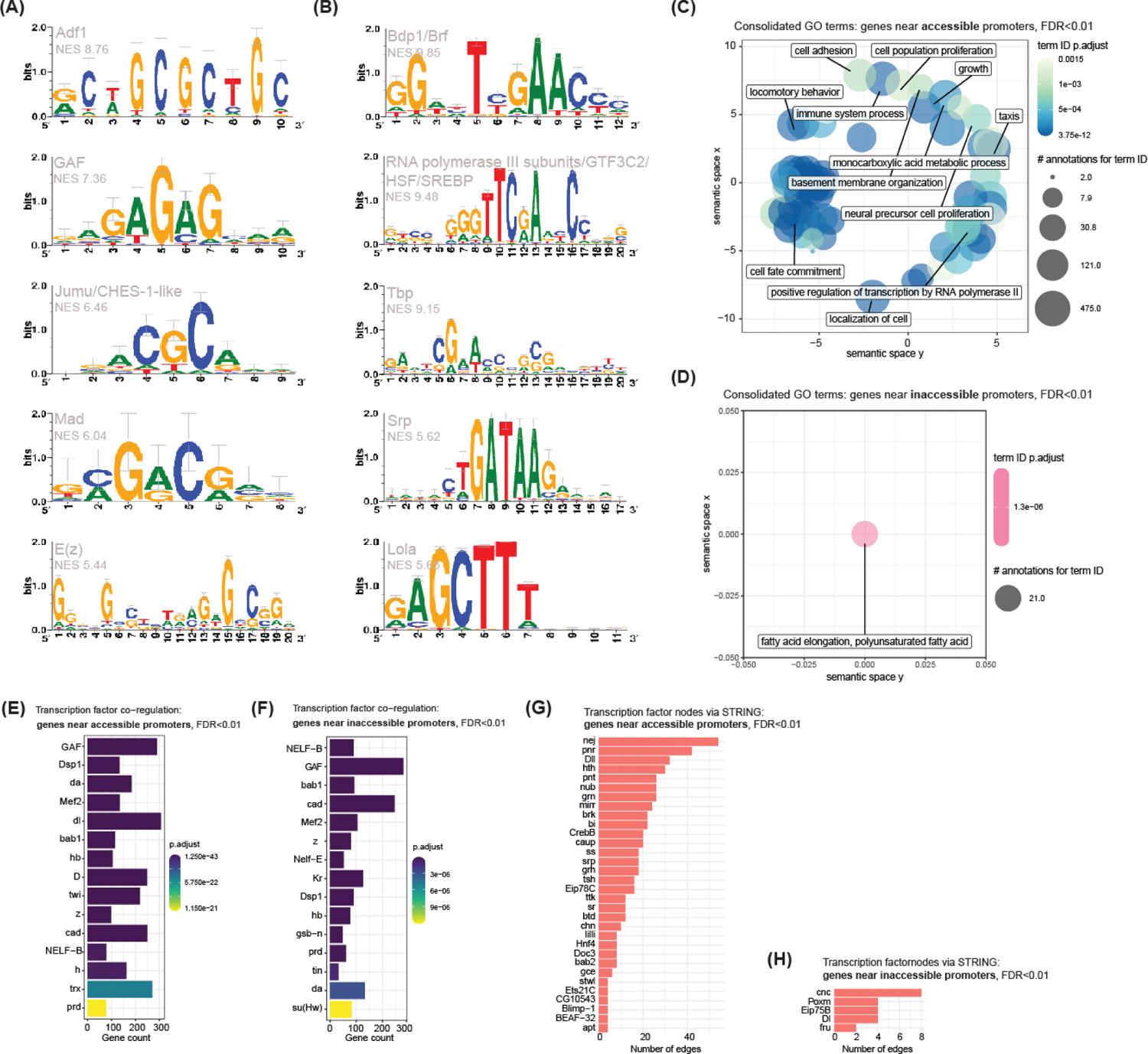
Promoter regions in diapause are structurally altered to facilitate changes in gene regulatory networks controlling cell fate, protein synthesis and energy metabolism. **(A)** Highly overrepresented DNA binding proteins whose binding motifs become accessible in diapause. All results from i-cisTarget were obtained under default parameters (NES threshold=3.0, minimum overlap=40%, AUC threshold=0.01, recovery curve threshold=5000). **(B)** Highly overrepresented DNA binding proteins whose binding motifs become inaccessible in diapause. **(C)** Functional categorization via Gene Ontology using the set of genes near accessible promoters (within ±3 kb). Categories were consolidated using Revigo according to functional similarity and statistical significance to generate clusters of related function (similarity cutoff value=0.5). Labeled terms are the most representative of the cluster to which they belong. **(D)** Functional categorization via Gene Ontology of genes near inaccessible promoters. Consolidated using Revigo (similarity cutoff value=0.5). **(E)** Top 15 most significant upstream regulatory proteins predicted from DroID co-regulation analysis using the set of genes near accessible promoters as input. Gene count is the number of input genes regulated by a given transcription factor. **(F)** Top 15 most significant upstream regulatory proteins predicted from DroID co-regulation analysis using the set of genes near inaccessible promoters as input. **(G)** Transcription factors derived from STRING interaction network analysis using the set of genes near accessible promoters as input (cutoff = 4 edges). A node is defined here as a protein with which one or more different proteins interact. Interacting proteins are edges. Transcription factors were identified from the set of all output nodes, which included non-transcription factors, by cross-referencing our data with transcription factor data from Flymine and Flybase. **(H)** Transcription factors derived from STRING interaction network analysis using the set of genes near inaccessible promoters as input.

The binding motif for another essential transcription factor, Adf-1, was also enriched in accessible loci (fig. 4A). Adf-1 regulates the expression of genes required for a variety of tasks, such as the synthesis of dopamine, normal dendrite development and adaptive plasticity (England et al., 1990; Timmerman et al., 2013). Changes to neuronal secretory identity and the dopamine synthesis pathway are strongly correlated with diapause (Andreatta et al., 2018; Karp, 2021; Nagy et al., 2019). Notably, Adf-1 also helps mediate transcriptional pausing through association with GAF (Timmerman et al., 2013).

Other motifs that become accessible in diapause are similarly relevant for development, cell proliferation and cell fate determination. Motifs were recovered that bind factors such as Jumu, caup, CHES-1-like and Mad, supporting the idea that chromatin in diapause is arranged to respond to these factors’ binding by promoting tissue reorganization (fig. 4A). In particular, accessibility of the Mad and Aef1 binding motifs further suggests changes to the developmental trajectory of the young adult fat body (Falb & Maniatis, 1992). Mad is a Dpp-responsive *Drosophila* Smad responsible for maintaining structural integrity of the fat body during development (Maduzia & Padgett, 1997; Hoshizaki, 2005). Aef-1 is a repressor that binds and inactivates fat body enhancers such as the ecdysone response element that drives production of fat body protein 1 (fbp1) (Brodu et al., 2001). Fbp1 is the receptor of larval serum proteins 1 and 2 (Burmester et al., 1999). During the beginning of pupal metamorphosis, a major point of developmental transition for the fat body, larval serum proteins are taken up by fat body cells through fbp1. The synthesis of fbp1 is also restricted to this stage. The accessibility of the Aef1 binding motif in diapause implies that transcription of fat body protein 1 is enabled, perhaps priming the tissue for end stage remodeling upon the exit from diapause. In sum, the collection of motifs that become accessible in the diapause fat body send a unified message of stalled or otherwise altered developmental trajectory. The complete list of accessible motifs is available (supplemental table S5).

Motif enrichment analysis of genomic loci that become inaccessible during diapause showed very strong representation for a single motif (5’-TTCGAACC-3’) which binds many factors related to the function or assembly of the RNA polymerase III complex (fig. 4B). These include general transcription factors such as the TATA binding protein (Tbp), components of the TFIIIB initiation factor (Brf and Bdp1), and subunits of the RNA polymerase III complex itself (Lobo et al., 1992; Huet & Sentenac, 1992; Liao et al., 2003). Although this motif does not represent the canonical Tbp binding sequence, many Tbp-associated factors (TAFs) are known to regulate transcription from all three major RNA polymerases (Akhtar & Veenstra, 2011). It is possible that Tbp binds this motif through association with TAFs. Heat shock factor 1 (HSF1) and sterol regulatory element binding proteins (SREBPs) use a similar motif. HSF1 binding correlates with TATA occupancy and acts downstream of GAF-induced polymerase pausing to release a subset of RNAPII at the promoters of stress-responsive genes (Duarte et al., 2016; Shopland et al., 1995). SREBPs are major transcriptional regulators of enzymes catalyzing the synthesis of various lipid species in response to insulin, which in flies is a function generally localized to the fat body (Dif et al., 2006; Seegmiller et al., 2002). Recently it was shown that SREBPs not only regulate fat storage, but alter cell fate depending on the rigidity of the extracellular matrix (Bertolio et al., 2019).

Of particular interest was the inaccessibility of the GATA motif, which was especially enriched in results from HOMER (up to 27.6% of input sequences, p<0.01) (fig. 4B, supplemental fig. S.4, supplemental table S4). In mammals, proteins that bind the GATA motif mediate hematopoiesis and organogenesis (Hasegawa & Shimizu, 2017; Molkentin, 2000; Whitcomb et al., 2020). Proteins that bind the GATA motif in *Drosophila* similarly regulate the division, specification and differentiation of precursor cells to the blood-analogue tissue hemolymph. Specifically, the GATA factor serpent (srp) is a critical driver of differentiation in the larval fat body (Hoshizaki, 2005). The inaccessibility of this motif in diapause may imply a tissue-specific modification of the development of this tissue. The complete list of inaccessible motifs is available (supplemental table S6). Together, results from our motif enrichment analysis suggest that the chromatin landscape in diapause is structured to facilitate transcriptional changes in lipid metabolism, protein synthesis, fat body development, cell fate and division.

Given that most differentially accessible loci mapped to promoters, we then considered that the transcriptome of the fat body in diapause could be modeled through examination of nearby genes. We performed functional categorization of genes that contained a differentially accessible promoter. Genes with promoters that become accessible in diapause are largely related to development, cell proliferation and differentiation, but also categorize into less frequent functions of organismal locomotion, RNAPII-mediated transcription, DNA replication and innate immunity (fig. 4C). Notably, these results agree with interpretations from our motif analysis that implicate development and protein synthesis as two processes in diapause differentially regulated through chromatin conformation. Due to the abundance of redundant GO terms produced from this dataset (mainly in the class of development), we leveraged the clustering algorithm from Revigo (Supek et al., 2011) to consolidate terms according to functional similarity and statistical significance. The full list of GO terms prior to consolidation is available (supplemental table S7).

Genes with promoters that become inaccessible in diapause grouped almost exclusively into categories related to fatty acid synthesis (supplemental table S8). As such, consolidation of terms resulted in a single cluster, of which the most representative term was elongation of polyunsaturated fatty acid (fig. 4D). This result suggests fatty acid synthesis is greatly altered through chromatin changes in diapause, perhaps to accommodate both the cuticular modifications and metabolic changes required for diapause survival and longevity.

Taken together, we find that genes regulated by differential promoter accessibility in diapause can be categorized into functions affecting development, fatty acid synthesis, regulation of RNA polymerase, and regulation of cell fate specification. These findings agree with our interpretations from motif analysis.

We also leveraged the Drosophila Interactions Database (Yu et al., 2008; Murali et al., 2011) to derive candidate upstream regulatory proteins from co-regulated genes with differentially accessible promoters. Like motif enrichment analysis, co-regulation analysis functions as a method to predict unifying drivers of transcription. However, whereas motif analysis returns predictions for DNA binding proteins, co-regulation analysis may capture additional modifiers of transcription that grant specificity to the genomic locations of DNA binding proteins. It is worthwhile to note that DroID is rather small (149 total transcription factors with interaction data from RedFly and ModENCODE). The database therefore may exclude some factors responsible for transcriptional regulation in diapause.

Encouragingly, we saw that many genes with differentially accessible promoters are co-regulated by GAF, or physical interactors of GAF, which corresponds with the significance of the GAGAG motif in diapause (fig. 4 E,F). GAF-associated factors that facilitate transcriptional pausing were also enriched in our results from DroID, including bab1 and subunits of NELF (Tsai et al., 2016). We noted similarity in results derived from genes whose promoters become accessible and those whose promoters become inaccessible. Broadly, the shared theme between datasets suggests stalled transcription and differential regulation of mesodermal development in the diapause fat body. Our results suggest this occurs through the reorganization of chromatin at promoter regions for genes interacting with master regulators such as daughterless (da), Mef2, dorsal (dl, an NF-κB factor), zeste (z) or diachete (D) (Smith & Cronmiller, 2001; Markstein et al., 2001; Kal et al., 2000). RNAPII stalling is a crucial mechanism of homeotic gene control in development (Levine, 2011; Core & Lis, 2009). It follows that genes with promoters containing the GAF motif, or motifs of GAF binding proteins, are more likely to be regulated through paused RNAPII, insofar as GAF recruits elements of the NELF complex. We also note the broader centrality of GAF in these results; GAF physically interacts with many developmental regulators, including both Polycomb group (PcG) proteins and Trithorax group (TrxG) proteins, as well as shared components of each (Horard et al., 2000; Schuettengruber et al., 2009; Salvaing et al., 2006; Déjardin & Cavalli, 2004; van Steensel et al., 2009; Brock & van Lohuizen, 2001). Together, PcG and TrxG proteins mediate transcriptional priming at the promoters of many genes in pluripotent cells (Dillon, 2012). These data support our earlier interpretation that chromatin in diapause is structured to modify development in the fat body upon diapause termination. Complete lists of candidates derived from this analysis are available (supplemental tables S9, S10).

To better understand the gene regulatory network suggested by our ATAC-seq data, we next sought to discover transcription factor nodes within the set of genes with altered promoter accessibility. These may represent important target genes for our candidates derived from DroID. We therefore used the STRING database collection (version 11.5) to generate interaction networks for genes with altered promoter accessibility. From the interaction networks generated, we extracted and filtered the nodes using data from Flymine (Lyne et al., 2007) and Flybase (Gramates et al., 2022) to arrive at datasets only including transcription factors. The most interactive transcription factors within the set of genes with accessible promoters were homeotic, reiterating interpretations from our previous analyses (fig. 4G). Nejire (nej) was our most interactive result. Nej is the *Drosophila* orthologue of CREB binding protein (CBP), a transcriptional coactivator that mediates many aspects of mesodermal development as a component of both histone acetyltransferase and methyltransferase complexes. Nej directly regulates the activity of signal transduction pathways such as Notch, hedgehog, dl, Dpp (TGF-β), and circadian rhythm, likely explaining its position as the most interactive node (Luo et al., 2017; Jia et al., 2015; Akimaru et al., 1997; Waltzer & Bienz, 1999; Lilja et al., 2003; Maurer et al., 2016). Indeed, most transcription factors extracted from these data are dependent on the activity of nej or are downstream within the pathways nej regulates. Nej also attenuates the activity of FoxO through acetylation, which further extends the network of modes by which nej controls cellular growth, proliferation and development (Brunet et al., 2004; Matsuzaki et al., 2005; Daitoku et al., 2004).

Many more transcription factors were obtained from the set of genes with accessible promoters than the set of genes with inaccessible promoters, in spite of the similarity in input size for each dataset (306 genes with accessible promoters, 348 genes with inaccessible promoters) (fig. 4H). This may be explained through the hypothesis that transcriptionally silenced genes in diapause are primarily downstream effectors, although the reason for this being the case is unknown. Of the few transcription factors derived, we found a common function in maintaining stem cell homeostasis. Cnc mediates the activation of stress resistance genes to promote cellular quiescence by managing the intracellular redox state (Pitoniak & Bohmann, 2015; Bayliak et al., 2020). Cnc deficiency is reported to delay morphogenesis and disrupt energy metabolism by lowering the rate of oxidative phosphorylation. Poxm, Eip75B (PPARY) and Delta (Dl) are also critical mediators of the balance between proliferation and quiescence. Notably, Eip75B mediates stem cell fate as a function of lipid metabolism, instantiating our results from GO (Zipper et al., 2020). The complete lists of genes with differentially accessible promoters, as well as their interaction data per gene, are available (supplemental tables S11, S12).

To refine our predictions, we then asked how well the upstream regulatory proteins predicted from motif analysis agreed with those predicted from co-regulation analysis. Considering the promiscuous nature of GAF, we expected many of the proteins predicted through DroID would function as GAF binding partners, rendering their activity dependent on GAF (Lomaev et al., 2017). This led us to a hypothesis wherein the differential accessibility of promoters with GAGAG motifs is determined by GAF-interacting proteins in a combinatorial fashion.

By mapping the physical interactions between our candidate upstream regulatory proteins, we found that, indeed, GAF functions as a hub for other factors predicted through DroID (fig. 5A). The significance of GAF in both datasets for accessible loci highlights the importance of chromatin remodeling in diapause and provides context to the molecular mechanisms by which these changes are mediated throughout the genome. Other factors shared between datasets for accessible loci, such as CtBP, E(z), grh, GATAe and z, have a common role in repression, particularly against genes responsible for the balance between stem cell maintenance and differentiation (Hoang et al., 2010; Aihara et al., 2013; Blastyák et al., 2006; Okumura et al., 2016).

**Figure 5:**
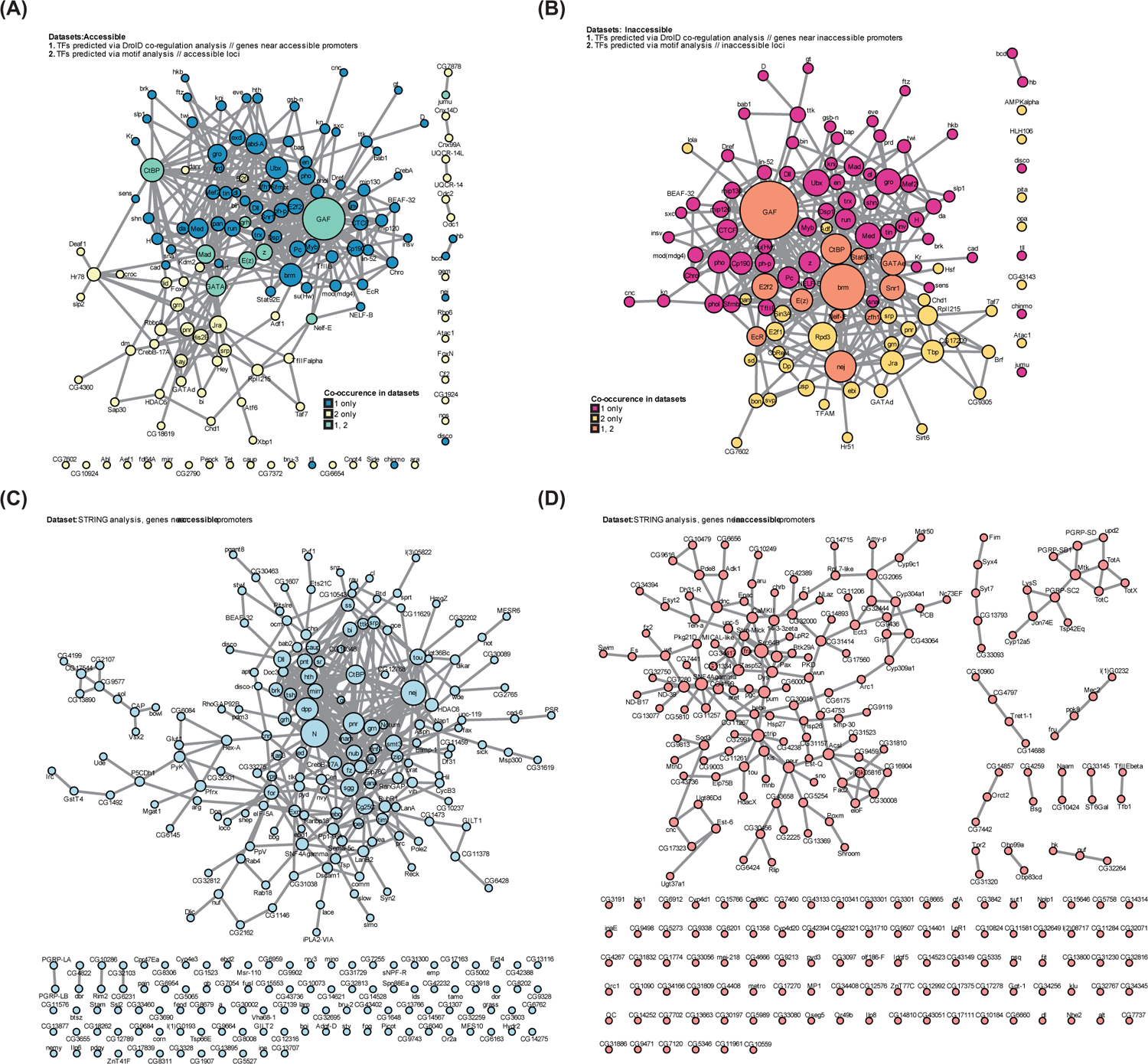
Identification of GAF and Brahma as key regulators of chromatin-mediated gene control in diapause, targeting Notch-interacting genes. **(A)** Interactions between upstream regulatory proteins predicted from DroID co-regulation analysis and motif enrichment analysis of accessible loci. Node size is determined by the number of the node’s edges found within either network. A confidence score of 0.4 and an FDR stringency of 0.05 were used to determine interactions through STRING. **(B)** Interactions between upstream regulatory proteins predicted from DroID co-regulation analysis and motif enrichment analysis of inaccessible loci. **(C)** Interactions within the set of genes near accessible promoters. **(D)** Interactions within the set of genes near inaccessible promoters.

Along with GAF, components of the nucleosome remodeling Brahma (SWI2/SNF2) complex were found to be highly interactive in both datasets for upstream regulatory factors derived from inaccessible loci (fig. 5B). Brahma facilitates chromatin remodeling necessary for developmental processes such as cell cycle progression and stem cell renewal (Shi et al., 2014; Jin et al., 2013). The inaccessibility of EcR-regulated promoters has important implications for diapause (Karp, 2021; Poupardin et al., 2015; Guo et al., 2021). Ecdysone signaling is a long-studied determinant of developmental progression versus delay in insects; the agreement of both datasets on this factor implies that ecdysone-mediated transcription may be downregulated in diapause to maintain the organism in a metamorphically static state. In addition, the agreement of datasets upon nej suggests that, while the promoter of nej itself is accessible, the promoters of its target genes are not. This is consistent with our model for diapause-associated genome priming, where the pro-development activity of nej will be permitted upon exit from diapause.

To discover gene regulatory networks outside transcription factor regulation, we also performed network analysis for all genes with differentially accessible promoters. From our previous results, we learned that nej was highly impactful among genes whose promoters become accessible in diapause. However, within the context of other genes in this set that do not encode transcription factors, we were able to discern that, of the many signal transduction pathways regulated by nej, Notch and Dpp appear to be the most highly supported (fig. 5C). Both pathways are essential to the maintenance of stem cell identity and the induction of cellular quiescence, depending on environmentally and developmentally timed cues (Dey et al., 2016; Firth et al., 2010; Sood et al., 2022). Interestingly, Notch signaling has also been shown control differentiation through its effects on fatty acid metabolism (Garcés et al., 1997; Jabs et al., 2018). These results provide mechanistic context to our most significant pathways identified through GO.

Corresponding with the low number of transcription factors derived from our set of genes with inaccessible promoters, we also found little cohesion around any one gene when non-transcription factors were included (fig. 5D). Considering this in light of our functional analysis of these genes, we did find general consensus of genes related to fatty acid synthesis, but also genes regulating the activity of Notch and cellular adhesion. The lack of interactions within this dataset may suggest the absence of specific transcription factor regulation, where transcription of silenced genes is instead regulated by more general factors responsible for chromatin decompaction. This interpretation agrees with our earlier finding that GAF and the Brahma remodeling complex co-regulate genes with inaccessible promoters.

### Summary of ATAC-seq results

To summarize our ATAC-seq analysis, we found that widespread changes to the chromatin landscape in diapause—particularly promoter regions—are likely dependent on GAF. In addition to its roles in recruiting both PcG and TrxG proteins, GAF may also recruit NELF to induce a poised state of RNAPII at the promoters of certain genes. Our data show that the genes affected by these changes are integral to the growth and development of the fat body; their poising by GAF and NELF could indicate the tissue is transcriptionally primed to resume development upon the detection of environmental cues signaling the termination of diapause. Indeed, we found further indication of stalled development through the differential accessibility of promoters for genes regulating the homeostatic balance of progenitor cells to maintain an undifferentiated, quiescent state. We also observed significant loss of chromatin accessibility around RNAPIII-related binding sites, as well as the promoters of genes supporting fatty acid synthesis. These results engage two more processes relevant for quiescent stem cell physiology, the reduction of protein synthesis and the modification of fatty acid metabolism. Lastly, we note the importance of Notch in our data; Notch is a master regulator of cellular quiescence and differentiation with the ability to affect fatty acid metabolism. Many genes of the Notch signaling pathway are regulated through chromatin changes in diapause, reinforcing the idea that the fat body may retain characteristics of an undifferentiated tissue in diapause, poised for the resumption of development. Together, our data from ATAC-seq support a model wherein diapause induces a poised state of chromatin, both to maintain cellular quiescence and to facilitate rapid, pro-development transcription upon diapause termination.

### Transcriptional changes in diapause support pathways that maintain cellular quiescence

Changes to chromatin accessibility do not necessarily imply corresponding changes in transcription, however, so we also performed RNA-seq on fat body tissue using replicate conditions from our ATAC-seq experimental design to address the hypothesis that the transcriptome supports those processes suggested by chromatin structural changes in diapause. RNA-seq libraries were sequenced to an average depth of approximately 52 million reads per sample and processed using a similar workflow as used for our ATAC-seq data. An average of 94.7% (±1.0% standard deviation) of reads mapped uniquely to the Flybase dm6 reference genome. Replicates clustered according to condition (supplemental fig. S.5). Differential expression analysis was conducted in DESeq2 using the same criteria as in ATAC-seq; genes were considered significantly upregulated in diapause if both diapause-inducing conditions showed significant upregulation (FDR<0.01) and either standard condition did not. Likewise, genes were considered significantly downregulated in diapause if standard conditions showed significant upregulation (FDR<0.01) and diapause-inducing conditions did not. Using this setup, our analysis returned 2,373 significantly upregulated genes and 1,989 significantly downregulated genes in diapause (fig. 6A). The diapause-associated changes to circadian gene expression—such as the upregulation of *period* and downregulation of *clock—* was observed through our analysis, suggesting our experimental design accurately represents diapause physiology (Salminen et al., 2015; Homma et al., 2022). In addition, JH-responsive genes Obp99a and Fad2 showed expected changes in transcription according to the presence or absence of S-methoprene, demonstrating the success of our supplementation approach (supplemental fig. S.6). The full list of differentially expressed genes is available (supplemental table S13).

**Figure 6:**
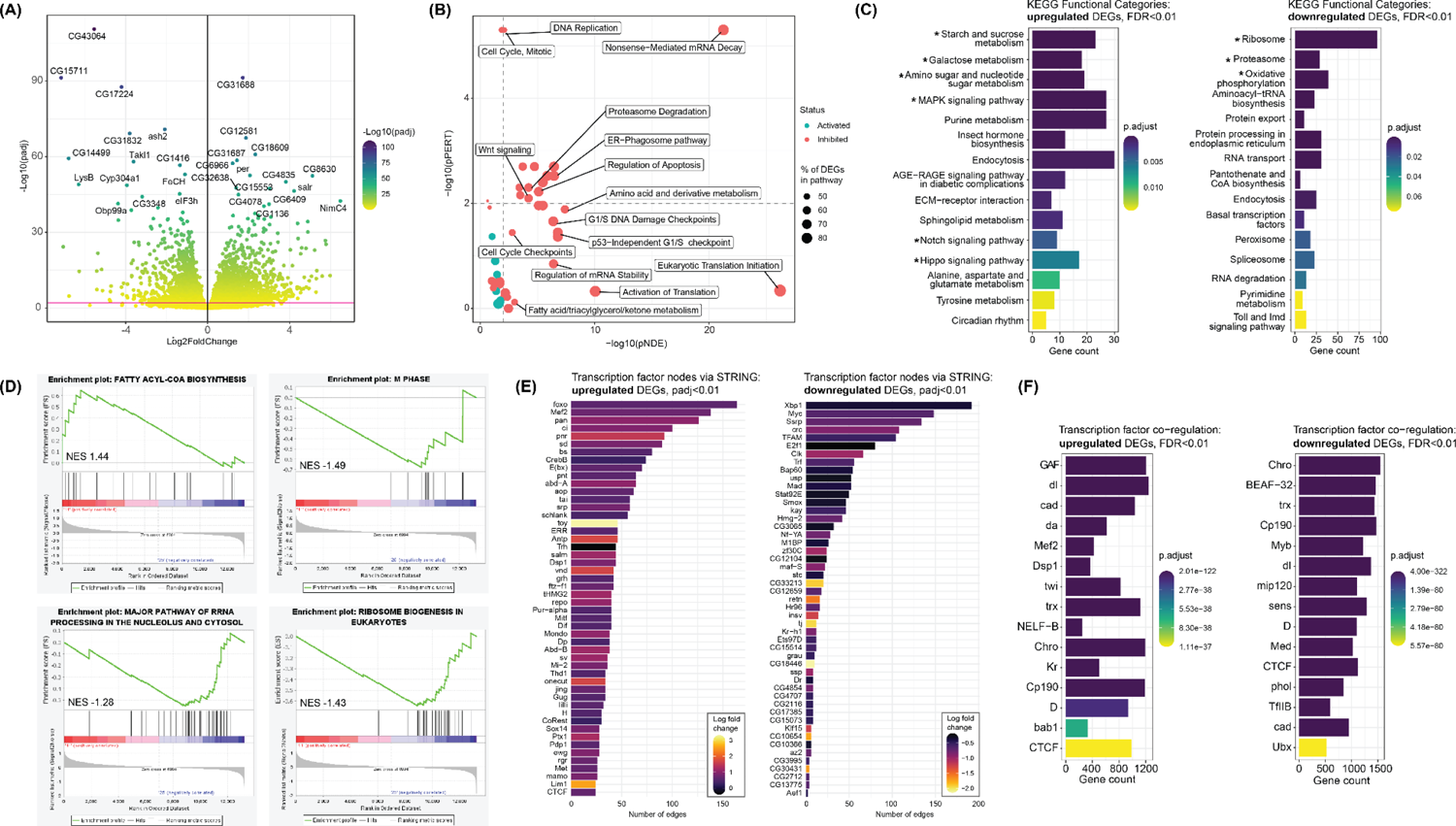
Diapause-associated transcription promotes the maintenance of cellular quiescence in the fat body. **(A)** Volcano plot showing the distribution of significance and log fold change for differentially expressed genes in diapause. FDR-adjusted p value (padj) significance threshold of padj<0.01 is indicated by the magenta line. **(B)** SPIA performed on all differentially expressed genes with padj<0.01 (DEGs). Dashed lines indicate significance thresholds of 0.01 for pPERT and pNDE. **(C)** KEGG pathway analysis performed separately for upregulated (left) and downregulated (right) DEGs. Starred terms indicate relevance to quiescent cell physiology mentioned within the main text. **(D)** GSEA performed on all DEGs. Gene matrices were derived from KEGG and Reactome to generate an unbiased set of query pathways. **(E)** Transcription factors derived from STRING interaction network analysis using the set of upregulated DEGs (left, cutoff = 25 edges), or downregulated DEGs (right, cutoff = 2 edges) as input. Complete lists of transcription factors (no edge cutoff) are available as supplements. Transcription factors were identified by cross-referencing our DEGs with transcription factor data from Flymine and Flybase. **(F)** Top 15 most significant upstream regulatory proteins predicted from DroID co-regulation analysis using the set of upregulated DEGs (left) or downregulated DEGs (right) as input. Gene count is the number of input genes regulated by a given transcription factor.

We conducted topology-based pathway analysis to understand the functional impact of differential transcription in the diapause fat body. Using Signaling Pathway Impact Analysis (SPIA) in conjunction with the Reactome (version 81) pathway database, we uncovered transcriptionally inhibited pathways controlling protein synthesis and degradation, as well as cell cycling and turnover (fig. 6B). We interpret the coincidence of these pathways to indicate maintenance of the existing cell population. We also observed significant transcriptional changes to fatty acid metabolism. Interestingly, although the analysis shown here was performed using differentially expressed genes with an FDR-adjusted p value cutoff of 0.01, the status of this pathway becomes activated under an FDR-adjusted p value cutoff of 0.05. This switch was exclusively observed for the three pathways directly related to fatty acids, which includes two others dependent on lipid signaling via PPARα. This bivalence may reflect diverging regulation of fatty acid synthesis and β-oxidation, which is a common theme among quiescent cells. The complete list of pathways derived from this analysis is available (supplemental tables S14, S15). In sum, our results from SPIA are consistent with our previous assertions derived from ATAC-seq functional analyses, where the diapause genome is structurally and transcriptionally altered to suppress cellular proliferation and protein synthesis, perhaps trading developmental progress for the maintenance of stem cell homeostasis.

To test the robustness of our results from SPIA, we performed additional pathway analyses using KEGG (Kyoto Encyclopedia of Genes and Genomes) pathway overrepresentation and Gene Set Enrichment Analysis (GSEA). The most significant KEGG pathways within the set of upregulated genes were metabolic (fig. 6C, left). This may represent a shift in energy source, reciprocating our previous observations of diapause-associated changes to fatty acid metabolism. The functional significance of MAPK signaling is less clear. The diversity of MAPK signaling outcomes ranges widely, and though the pathway is often associated with proliferation, downstream effectors of MAPK signaling can also inhibit the cell cycle and maintain a primed state of pluripotency (Shilo, 2014; Nir et al., 2012; Dundes & Loh, 2020). KEGG pathway analysis also corroborates our results from SPIA by showing significant upregulation of Notch and Hippo signaling, two pathways controlling the balance between cellular quiescence and activation. Hippo signaling coordinates the restriction of growth during development and the maintenance of stem cell quiescence in *Drosophila* (Halder & Johnson, 2011; Staley & Irvine, 2012; Ding et al., 2016; Jin et al., 2013). As previously addressed, Notch signaling also plays an important role in maintaining *Drosophila* stem cell quiescence (Bjornson et al., 2013; Bernard et al., 2010; Sood et al., 2022; Anant et al., 1998). Its reappearance in our RNA-seq reaffirms the importance of this pathway for diapause, well as the robustness of our data.

Meanwhile, KEGG pathway analysis of downregulated genes revealed an overwhelming representation of the ribosome, followed by the proteasome, further validating results from SPIA (fig. 6C, right). The loss of oxidative phosphorylation is a typical feature characterizing primed pluripotency (Shyh-Chang & Ng, 2017). This interpretation is additionally supported in context of the highly upregulated pathways for sugar and starch metabolism, together suggesting a shifted energetic preference towards glycolysis, which often accompanies quiescent cell metabolism along with changes to fatty acid metabolism.

Results from GSEA corroborated our findings from previous pathway analyses, where we observed significant downregulation of cell cycle-related transcription (M phase NES = −1.49) and protein synthesis (major pathway of rRNA processing in the nucleolus and cytosol NES = −1.28, ribosome biogenesis in eukaryotes NES = −1.43) (fig. 6D). As in our SPIA results, we also found altered regulation of fatty acid metabolism (fatty acyl-coA biosynthesis NES = 1.44), though the directionality of this result was less certain. Whereas other results from GSEA showed a clear leading edge subset of significant transcripts, there was much greater spread in the transcripts corresponding to fatty acyl-coA biosynthesis between positive and negative correlation. This reinforces the ambiguity, but also the significance, of transcriptional changes to fatty acid metabolism in diapause that were determined through SPIA.

We then sought to unify these changes by searching for highly influential transcription factors within our differentially expressed genes. We first used the interaction network-based method previously described for genes with differentially accessible promoters and identified many transcriptionally upregulated factors directing development and cell fate specification (fig. 6E, left). Many of these factors have bivalent roles in promoting and repressing the differentiation of progenitor cells, depending on cues delivered to the stem niche (Tsuji et al., 2008; Sinenko et al., 2009). FoxO, the most interactive transcription factor of this set, is a well characterized regulator of many aspects of physiology, including apoptosis, gluconeogenesis, cell cycle arrest, redox homeostasis, differentiation, stem cell maintenance and longevity (Accili & Arden, 2004; Zhang et al., 2011; Chen et al., 2019; Jünger et al., 2003). FoxO is widely considered to have both pro-longevity and tumor suppressive effects through its regulation of target genes such as *cycE* (cyclin E), *lola, nej* or *brummer* lipase, which together skew cellular metabolism towards tissue maintenance and FAO as an energetic adaptation (Birnbaum et al., 2019; Molaei et al., 2019). Concerning the possibly competing actions of FoxO and those highly interactive factors promoting development within this dataset, one hypothesis may be that although these factors are transcriptionally upregulated, they remain untranslated. This agrees with our data that suggests protein synthesis is downregulated in diapause, as well as the idea that diapause induces a state of priming. Translational repression is a common feature of replicative dormancy that can be achieved through the repression of translation initiation factors, as well as the preservation of certain mRNAs in a manner dependent on poly-A tail structure (Marescal & Cheeseman, 2020; Passmore & Coller, 2021).

Correspondingly, insofar as FoxO is known to oppose the actions of Xbp1 and Myc, these were the most highly interactive of all downregulated transcription factors (fig. 6E, right) (Zhou et al., 2011). Myc mediates myriad processes instructing cellular growth, including ribogenesis and lipogenesis, two processes our data suggest are downregulated in adult diapause (Grifoni & Bellosta, 2015). Changes to the expression of Myc in the fat body alone are sufficient to drive systemic effects on growth; these changes can be induced through upstream alterations to ecdysone or insulin/insulin-like (IIS) signaling, which are also pathways known to mediate the entrance or maintenance of diapause (Delanoue et al., 2010; Hutfilz, 2022). Other highly interactive transcription factors derived from the set of downregulated genes include Bap60, Ssrp and crc, three factors responsible for facilitating Hox gene expression through chromatin remodeling and ecdysone signaling, respectively (Shi et al., 2014; Möller et al., 2005). Along with Xbp1, crc also functions during the unfolded protein response expected during stress or development to regulate glycolytic gene expression. Trf enables ribogenesis as a component of the RNA polymerase III complex, underscoring our interpretation from ATAC-seq that protein synthesis is inhibited. E2F1, the transcriptional regulator of cyclin E, was also significantly downregulated in this dataset, though its fold change is small (Denechaud et al., 2017). *cycE* itself was significantly downregulated (supplemental fig. S.6). Cyclin E drives the mitotic entry into S phase, and so the downregulation of E2F1 may support our interpretation that diapause causes temporary exit from the cell cycle. Complete lists of transcription factors derived from RNA-seq analysis are available (supplemental table S16.)

We also used DroID co-regulation analysis to uncover transcription factors responsible for orchestrating the diapause transcriptome. As in our set of genes with accessible promoter regions, GAF was identified as the most significant factor predicted to regulate the expression of upregulated genes (fig. 6F, left). This result is reflective of the centrality of GAF as a binding partner for many transcription factors driving development, as well as its role recruiting chromatin remodeling factors for transcriptional priming. Along with GAF, we found other TrxG members such as z and trx. However, we also note that the TrxG member *ash2* is among our most significantly downregulated genes, inviting questions about the regulatory mechanisms used in diapause to reorganize chromatin (supplemental fig. S.6). Other master regulators of development were found to coordinate upregulated gene expression in diapause, such as da, which, among its many targets, promotes the expression of the CDK inhibitor Dacapo (Dap) (Sukhanova & Du, 2008). The high significance of Mef2, a major effector of MAPK signaling, may explain the pathway’s upregulation in our KEGG pathway analysis (Blanchard et al., 2010). Twist (twi) is an important factor in the Notch-dependent negative regulation of differentiation, repeating a theme present in all results from our RNA-seq (Anant et al., 1998).

DroID co-regulation analysis of downregulated genes showed an abundance of highly significant insulator proteins including CTCF, CP190, BEAF-32, Chro, mod(mdg4), and the dREAM repressive complex subunits Myb, E2F2, lin-52, mip120 and mip130 (fig. 6F, right) (Bohla et al., 2014; Santana et al., 2020). The reason for this might be connected to the strong downregulation of ribosomal RNA and tRNA-producing genes observed in our functional analyses. Unoccupied RNAPIII promoters can function as barrier elements to mediate insulation against spreading heterochromatin formation (Raab & Kamakaka, 2010; Valenzuela et al., 2009). It follows that the high representation of insulator proteins co-regulating downregulated genes could indicate the use of these genes not for transcription, but as a means to poise chromatin in a specific conformation. Interestingly, Hox promoters loaded with paused polymerases also exhibit intrinsic insulator activity (Chopra et al., 2009). This could imply a dual role of the diapause chromatin landscape for priming post-diapause transcription and also maintaining cellular quiescence by limiting rRNA and tRNA transcription. Complete lists of candidates derived from co-regulation analysis of differentially expressed genes are available (supplemental tables S17, S18).

We sought to better understand the hierarchy of gene regulatory networks controlling diapause through a comparison of our differentially expressed transcription factors with those predicted to regulate their expression upstream. Interaction network analysis of upregulated genes showed high interaction with Hox genes such as *abd-A, abd-B* and *Antp*, which are regulated by two factors shared between datasets, Dsp1 and Mef2 (fig. 7A) (Decoville et al., 2001). Mef2 controls mesodermal development and circadian rhythm, as well as the fat body metabolic switch between energetic conservation and utilization (Blanchard et al., 2010; Potthoff & Olson, 2007; Azeez et al., 2014). These functions are achieved in a manner dependent on signals from the environment, highlighting the relevance of Mef2 in diapause. Dsp1 is a recruiter of PcG and TrxG (Déjardin et al., 2005). Together with GAF and CTCF, Dsp1 may represent a key player in the formation of the diapause-associated chromatin landscape.

**Figure 7:**
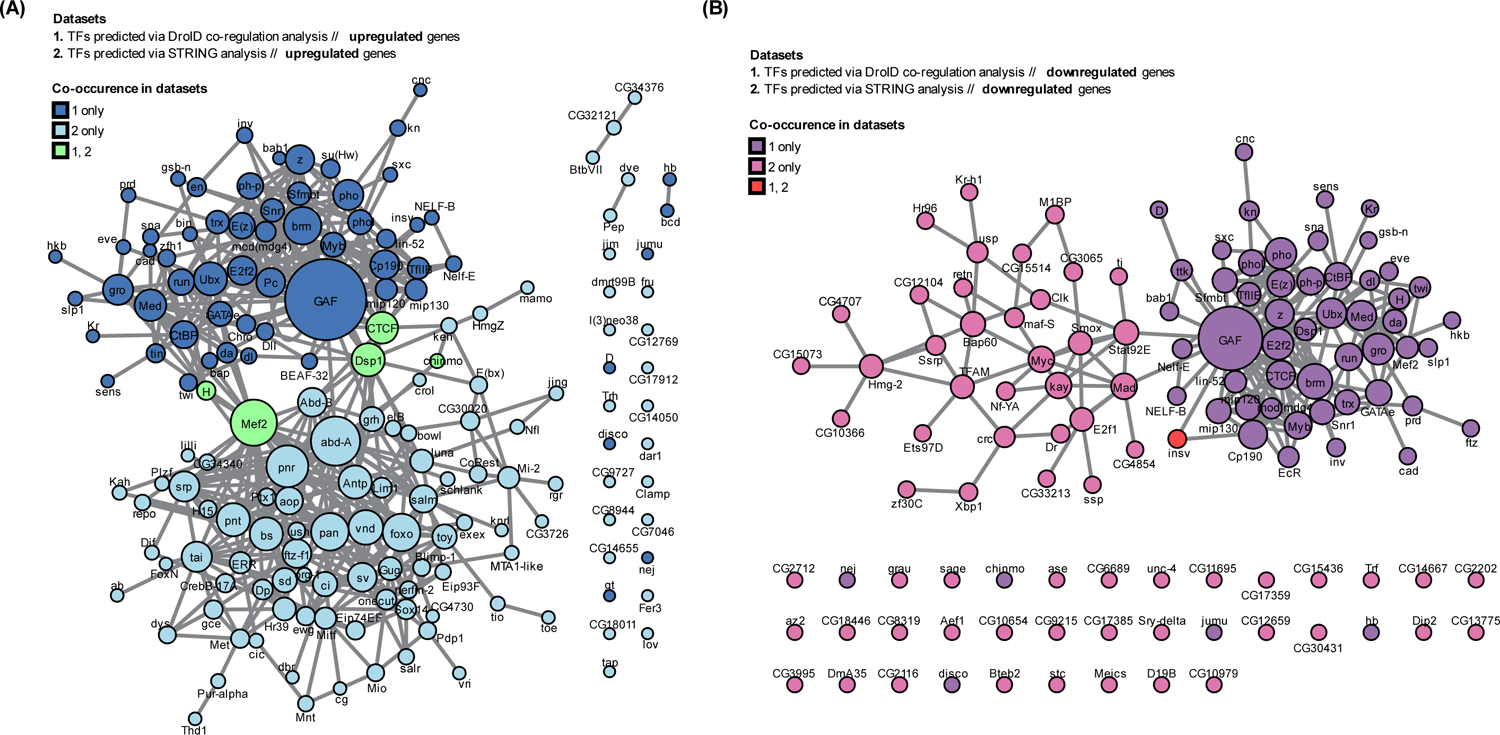
Identification of Mef2, Dsp1, insulators and hox genes as key regulators of transcription in diapause. **(A)** Interactions between upstream regulatory proteins predicted from DroID co-regulation analysis of upregulated DEGs and transcription factors identified through STRING interaction network analysis of upregulated DEGs. Node size is determined by the number of the node’s edges found within either network. A confidence score of 0.4 and an FDR stringency of 0.05 were used to determine interactions through STRING. **(B)** Interactions between upstream regulatory proteins predicted from DroID co-regulation analysis of downregulated DEGs and transcription factors identified through STRING interaction network analysis of downregulated DEGs.

We found very little overlap or interaction between datasets for downregulated transcription factors and those predicted to control their expression upstream (fig. 7B). This may be caused by the fact that our most significant upstream regulators were insulator proteins that likely all regulate our most significantly downregulated pathway, ribosome metabolism, through chromatin dynamics alone. This mode of transcriptional regulation is independent of transcriptional repressors, and so it may be the case that the downregulation of transcription factors we observe, such as Xbp1, is responsible for inhibiting pathways other than ribogenesis. These pathways could be those we determined through GO with lower gene counts than ribosome-related terms. GAF also interacts with insulator proteins, and so although it was not among our most significant co-regulators for downregulated genes, it is shown to be highly interactive (Melnikova et al., 2004).

### Chromatin structure and gene expression cooperate in parallel ways to prime the fat body for diapause termination

To understand the contribution of chromatin structure to transcriptional changes in diapause, we then compared our differentially accessible loci to the genomic loci of differentially expressed genes. Although we observed little overlap between categories (10.45% of differentially expressed genes matching genes that mapped to a differentially accessible locus), this is somewhat expected in consideration of the fact that the total number of genes with differentially accessible chromatin is 654. The highest possible overlap is therefore capped at 15%. In other words, using an FDR cutoff of 0.01, 69.7% of all genes with differentially accessible chromatin were also differentially expressed (fig. 8A). When we examined the directionality of these genes’ expression, we found a clear correlation between transcriptional upregulation and chromatin accessibility (fig. 8B). The prominence of GAF in our data for both accessible loci and upregulated genes may explain this result. This was not the case for differentially expressed genes within inaccessible loci, however; we found no correlation between inaccessible chromatin and the directionality of gene expression. This indicates that in diapause, changes to chromatin accessibility are more important for increasing transcription than decreasing. We also found no correlation between the fold change in chromatin accessibility and the fold change in gene expression for genes within differentially accessible loci (supplemental fig. S.7).

**Figure 8:**
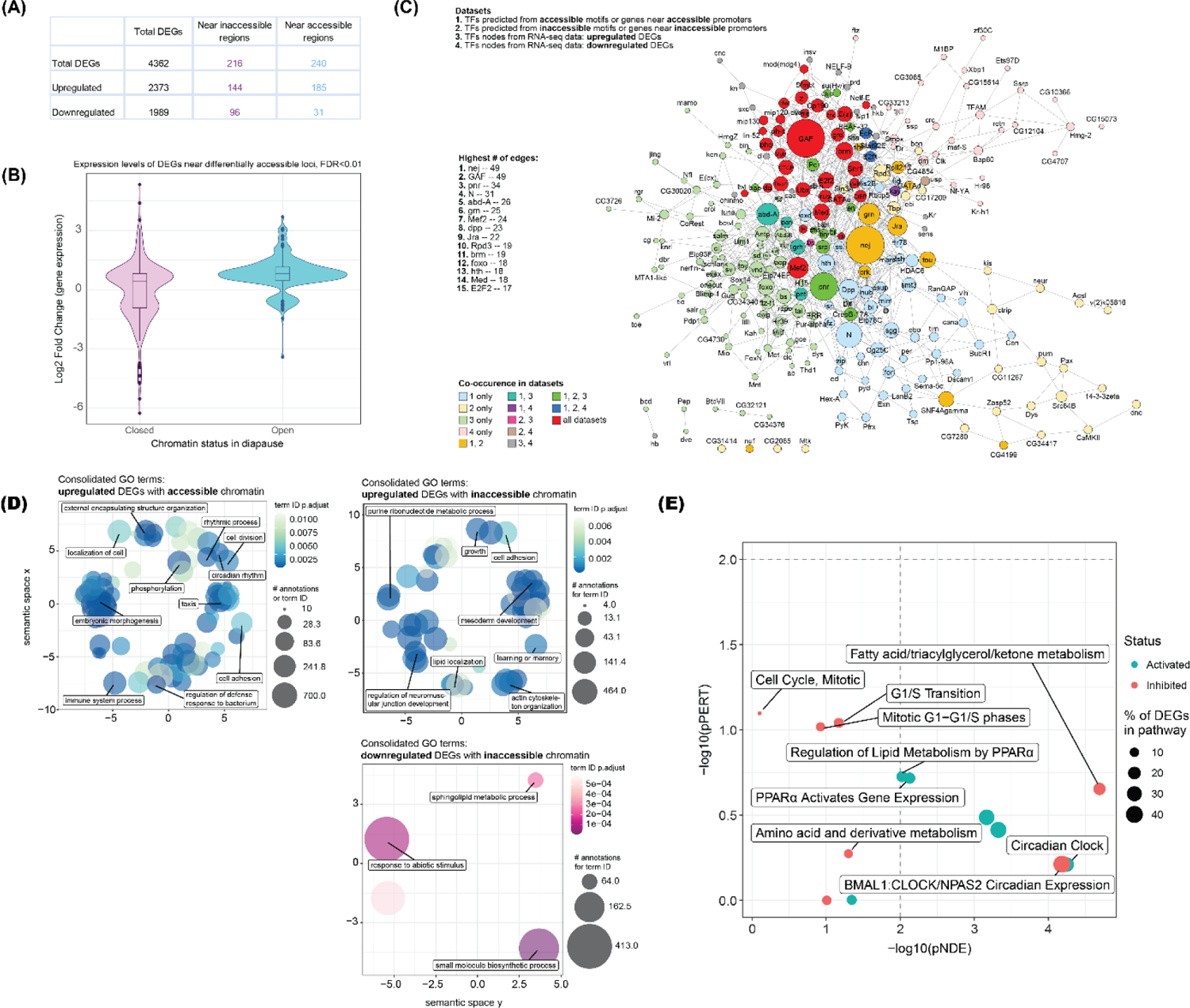
Chromatin structure cooperates with gene expression to promote cellular quiescence in diapause. **(A)** Quantification of genes with differentially accessible chromatin anywhere along the gene body that were also differentially expressed. **(B)** Violin plots comparing the directionality of chromatin accessibility (accessible or inaccessible) with the directionality of gene expression (upregulated or downregulated) for DEGs with differentially accessible chromatin. **(C)** Interactions between upstream regulatory proteins predicted from ATAC-seq and RNA-seq datasets. For ATAC-seq datasets (1 and 2), candidate regulatory proteins from motif analysis, co-regulation analysis and STRING interaction network analysis were combined according to whether they were derived from accessible or inaccessible loci. For RNA-seq datasets (3 and 4), candidate regulatory proteins from co-regulation analysis and STRING interaction network analysis were combined according to whether they were derived from upregulated or downregulated DEGs. Node size is determined by the number of the node’s edges found within either network. A confidence score of 0.4 and an FDR stringency of 0.05 were used to determine interactions through STRING. **(D)** Functional categorization via Gene Ontology using the set of upregulated DEGs with accessible chromatin (upper left), upregulated DEGs with inaccessible chromatin (upper right), and downregulated DEGs with inaccessible chromatin (lower right). There were too few downregulated DEGs of known functions with accessible chromatin to generate statistically significant GO terms. Categories were consolidated using Revigo according to functional similarity and statistical significance to generate clusters of related function (similarity cutoff value=0.5). Labeled terms are the most representative of the cluster to which they belong. **(D)** SPIA performed on all DEGs with differentially accessible chromatin. Dashed lines indicate significance thresholds of 0.01 for pPERT and pNDE.

Next, we compared our transcription factor predictions across datasets to arrive at a consensus set of transcription factors that may coordinate gene expression and chromatin reorganization in diapause. We found overrepresentation of highly interactive developmental drivers such as Ubx and EcR, as well as many components of chromatin reorganization complexes like Polycomb, Trithorax, Brahma, and the cell cycle inhibitory dREAM complex, which functions to maintain quiescence (fig. 8C). We also found high representation of all five GATA factors in *Drosophila* necessary for proper fat body and hemocyte development (srp, GATAe, GATAd, grain (grn) and pannier (pnr)) (Herranz & Morata, 2001). Pnr acts downstream of Stat and upstream of Mef2 and tin, three other highly represented factors, to control mesodermal cell fate specification through its role in Dpp signaling (Minakhina et al., 2011; Lovato et al., 2015; Gajewski et al., 1999). Pnr is also responsible for melanization in some insects, another process differentially regulated in diapause (Ando et al., 2018). Together, these results are consistent with the model wherein diapause induces changes in chromatin structure to allow for the pausing of development, the maintenance of stem cell quiescence, and genomic priming for post-diapause transcription.

Among the set of differentially expressed genes with affected chromatin, we found functional convergence of upregulated genes on related processes such as mesodermal development, cellular localization and adhesion. This appears to be independent of chromatin status, however, considering the similarity in results from differentially expressed genes with accessible or inaccessible chromatin. (fig. 8D). This observation could be explained if the primary role of chromatin conformational changes in diapause is to prime transcription for post-diapause life, rather than to direct transcription during diapause maintenance.

Downregulated genes associated with inaccessible chromatin were functionally related to fatty acid metabolism and the response to environmental stimuli. We were unable to functionally analyze downregulated genes associated with accessible chromatin because there were too few genes in this set with known functions. Complete lists of GO terms prior to consolidation via Revigo are available (supplemental table S19).

We also performed SPIA to validate our interpretations from GO analysis of differentially expressed genes with affected chromatin. Similar processes were recovered, though we note the conflicting directionality of pathways related to fatty acid metabolism and circadian rhythm (fig. 8E). This is reflective of the bivalence in fatty acid processes observed through our RNA-seq, and may suggest a more nuanced adaptation of these pathways in diapause, rather than a complete reversal of normal function. The complete lists of results from SPIA are available (supplemental table S20).

### The chromatin conformation of promoter regions in diapause resembles that of juvenile flies

Our data collectively suggest that at least a subset of fat body cells is maintained in a quiescent, undifferentiated state during diapause. During normal development, the larval fat body finishes development near the time of eclosion, when it undergoes histolysis and is replaced by new, terminally differentiated cells. During diapause, however, we propose that the larval fat is instead retained to some extent, and the progenitor cells resident within the adult fat body enter quiescence for the purpose of finishing development once conditions become favorable. This model implies the fat body in diapause is a developmentally incomplete tissue. To test this idea, we compared our ATAC-seq data with publicly available ChIP-seq data from Drosophila in the late larval stage. Promoter regions that become accessible in diapause aligned strongly with larval chromatin profiles for H3K9ac and H3K4me3, two markers of active or accessible promoters (fig. 9). Likewise, promoters that become inaccessible in diapause aligned very poorly. These results are consistent with our hypothesis that the fat body in adult diapause is stalled in a developmentally juvenile state.

**Figure 9:**
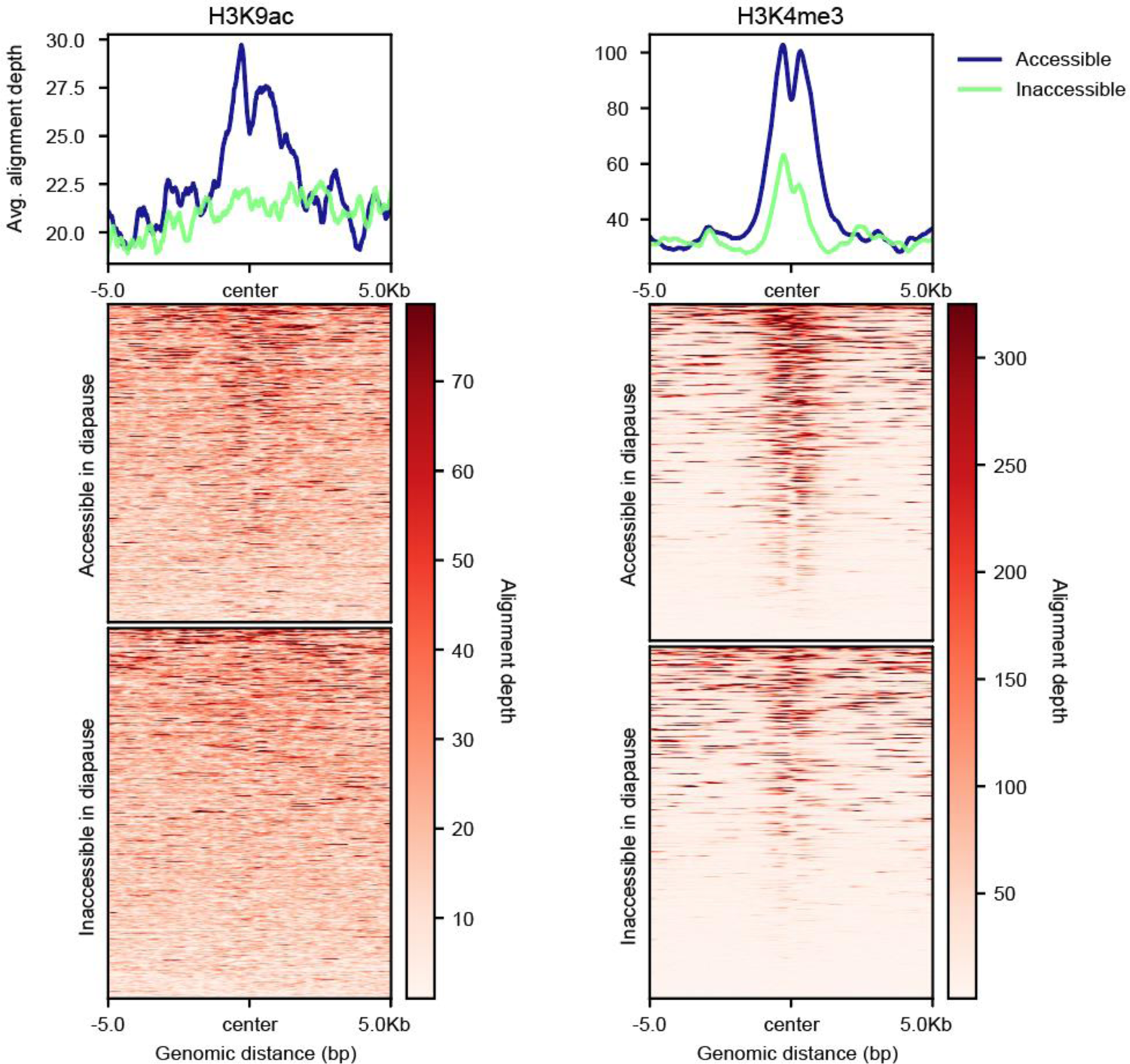
Promoter accessibility in the diapause fat body resembles that of juvenile flies. **(A)** Quantification of overlap between differentially accessible promoters in diapause and genomic loci significantly (FDR<0.01) bound to the accessible promoter mark H3K9ac in *Drosophila melanogaster* L3 larvae (modENCODE dataset ID #5102). Each row of the heatmap is a differentially accessible locus, binned into blocks of 50 bp and aligned at their centers. Alignment depth is the enrichment of larval H3K9ac at each bin. The profile plot above is the average score for each bin across loci. **(B)** Quantification of overlap between differentially accessible chromatin in diapause and genomic loci significantly (FDR<0.01) bound to the accessible promoter mark H3K4me3 in *Drosophila melanogaster* L3 larvae (modENCODE dataset ID #5097).

### Evidence for quiescent progenitor cells in the diapause fat body

The final step in fat body maturation is well-documented to entail histolysis of the larval fat and gradual replacement of this tissue by progenitor cell proliferation and differentiation. However, little effort has been dedicated thus far to the identification of this progenitor population. We next sought to locate quiescent progenitor cells in the fat body during diapause. Immunofluorescent tagging of the Notch ligand Delta was combined with an EdU (5-ethynyl-2’-deoxyuridine) incorporation assay to visualize stem cell proliferation over time in the diapause and post-diapause fat body. We found no clear EdU incorporation in fat body-derived Delta-positive cells from age-matched adults kept in standard conditions, nor from adults kept in diapause-inducing conditions (fig. 10A). This is consistent with the idea that fat body progenitor cells remain quiescent under these conditions. However, we were able to identify several Delta-positive cells with EdU-labeled nuclei in timepoints following the removal of flies from diapause-inducing conditions, though these observations were rare (fig. 10B). We interpret these results to support the possibility that diapause maintains a subset of fat body progenitor cells in quiescence during diapause.

**Figure 10:**
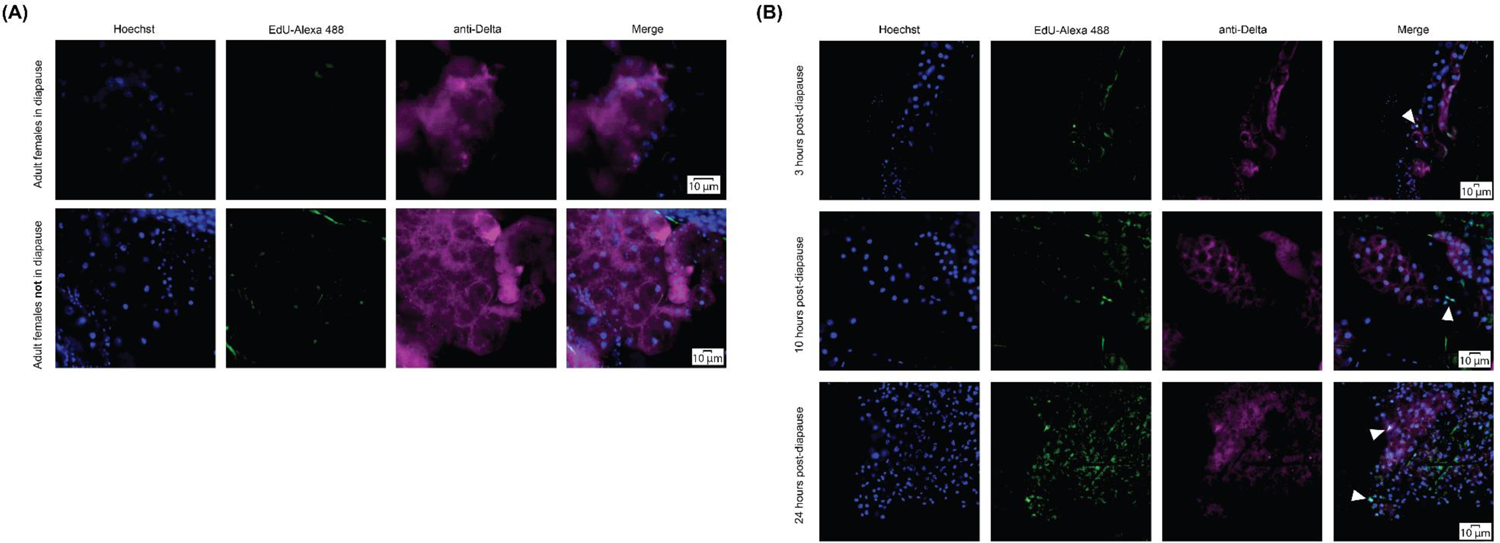
DNA synthesis is observed in progenitor cells of the fat body following exit from diapause, but not during diapause. **(A)** Representative images of EdU incorporation in Delta-positive cells of the fat body, dissected from 10-day-old w^1118^ virgin females kept in diapause-inducing conditions or standard conditions. **(B)** Representative images of EdU incorporation in Delta-positive cells of the fat body, dissected in time course post-diapause. White arrowheads indicate nuclei with visually distinct incorporation of EdU.

## Discussion

This work constitutes the first multiomic, tissue-specific and JH-controlled analysis of adult *Drosophila* diapause, as well as the first depiction of chromatin accessibility in the diapause fat body. These data will serve as a resource for future research in diapause, as well as related fields of development, agriculture, thermal tolerance and aging. In addition, our work introduces a model connecting progenitor cell quiescence to diapause-associated longevity. We found abundant hallmarks of quiescent cell physiology within our sequencing data, including the differential accessibility of homeotic genes, altered transcriptional regulation for fatty acid metabolism, the upregulation of FoxO, Notch, and Hippo-related transcription, downregulation of cell cycle, oxidative phosphorylation and ribosome biogenesis genes, and evidence for RNAPII pausing at the promoters of genes regulated by major transcription factors in development. These observations were supported *in situ* through the identification of replicative Delta-positive cells exclusive to timepoints post-diapause. We unify our findings through a hypothesis wherein, during diapause, progenitor cells in the fat body undergo cell cycle arrest. This may be mediated by global chromatin remodeling through the dREAM complex and GAF. In a context-dependent manner, GAF could recruit members of PcG, TrxG and the NELF complex to poise chromatin in a state that facilitates the rapid transcription of genes necessary for cell cycle reentry and resumption of development that accompanies diapause termination. The transcription that occurs during diapause in the fat body generally supports the maintenance of quiescent cell physiology and chromatin poising.

Given that our data are derived from the fat bodies of adult flies, however, from where do these progenitor cells originate? Two possibilities exist; under one assumption, the induction of diapause entails a reprogramming of differentiated cells in the adult fat body to a more stem-like state. More likely, perhaps, progenitor cells within the adult fat body are simply retained past the age they would normally undergo terminal differentiation. In our experiments, flies were placed into diapause-inducing conditions within 8 hours post-eclosion. It is known that the predecessor tissue to the adult fat body, the larval fat, completes histolysis in the span of about four days after eclosion, with most tissue death occurring in the first 2 days (Rizki, 1978; Hoshizaki, 2005). No literature has yet directly described the timeline (or existence) of those progenitor cells assumed to replace the larval fat through proliferation, but inferring from the fact that the young adult maintains a consistent presence of adipose throughout this process, we propose that diapause imposes the cessation of both larval fat histolysis and progenitor cell differentiation. The fat body would thus become a hybrid, developmentally stalled tissue in diapause. Both or either of the retained juvenile cell populations could account for our promoter accessibility data’s similarity to that of third instar larvae.

Hemocyte precursors may alternatively or additionally represent another fat body-resident population contributing to our data’s overwhelming quiescent signature. This idea is justified through our observation that Srp, a hemocytic marker, is transcriptionally upregulated while its GATA binding site is highly accessible. Hematopoietic niches in the adult are found embedded within the fat body and our sampling method for this tissue likely includes this population (Rehorn et al., 1996; Boulet et al., 2021). Cellular quiescence of hemocyte precursors could help to explain the lack of cell turnover our data suggests is the case during diapause: reduced macrophage production may present as cell cycle exit while simultaneously limiting the clearage of existing cells that accumulate damage under normal conditions. Indeed, under a model where somatic tissue is better maintained during diapause through the fat body-dependent reconfiguration of the insulin profile, cell clearance may become a superfluous expenditure of energetic resources.

Extrapolating from our model, the lifespan extension observed in diapause may be attributable to fat body quiescence. The fat body is a central mediator between organismal growth and environmental suitability (Koyama et al., 2020). Mitogenic signals from the fat body are required for other tissues to exit quiescence during development (Britton & Edgar, 1998; Sousa-Nunes, 2011; Hoshizaki, 2005). It follows that, upon the induction of quiescence, fat body-derived signals such as the Methuselah ligand Stunted, or the inaptly named growth-blocking peptides (GBP1, GBP2), may no longer be transmitted, leaving tissues outside the fat body incapable of growth, division and turnover to the extent they were under normal circumstances (Meschi et al., 2019; Koyama & Mirth, 2016). Insofar as the fat body is also a major regulator of systemic IIS and the immune system, loss of mitogenic signaling may in turn promote a widespread tissue maintenance program. This could be managed through metabolic adaptation (autophagy, or the downregulation of aerobic metabolism, as two possibilities), as well as the stimulation of plasmocytes responsible for disposing of hemolymph pathogens and supporting stem cell niche integrity through the secretion of extracellular matrix proteins (Gold & Brückner, 2015).

Distinguishing the precise cell population(s) responsible for our data’s quiescent signature will generate insight into the physiology of diapause-associated lifespan plasticity by identifying which cell type-specific signals may become altered through increased or decreased representation. With the installation of single-cell data for the adult fat body in Fly Cell Atlas, it may be possible to contextualize our results in terms of cell type contribution (Li et al., 2022). However, these data are currently only available for flies of mixed sex at three days of adulthood. Early studies in fat body morphology indicate larval adipocytes are entirely replaced by adult fat around day four post-eclosion (Rizki, 1978; Hoshizaki, 2005). More detailed knowledge of the fat body composition at all life stages would therefore benefit our understanding of how this tissue changes in diapause, and by extension, how its composition affects lifespan. For example, the relative abundance of larval adipocytes and fat body-resident immune cells may change in the diapausing adult when compared to age-matched controls. Bulk sequencing deconvolution tools such as CIBERSORTX could be used to better understand the cell composition represented by our data, under the hypothesis that adult diapause facilitates the retention of hematopoietic precursors, larval adipocytes and the adult fat body progenitor cells destined to replace them (Newman et al., 2019).

### MEF2 as a key regulator of diapause

In addition to GAF, we also highlight the possibility of a central role for Mef2 in our work. Mef2 is a multifunctional transcriptional activator whose specificity is guided by a plurality of context-dependent co-factors. First described for its role directing myogenesis, Mef2 is now known to broadly regulate the intersection of cellular proliferation, metabolic homeostasis, stress response and tissue remodeling in a variety of cell types (Potthoff & Olson, 2007). Of particular importance to this work, Mef2 controls lipid composition and lifespan in a manner dependent on external conditions (Zhao et al., 2020), as well as the expression of several genes within pathways regulating cell fate and quiescence, such as Notch (Caine et al., 2014). Accordingly, many differentially regulated signaling pathways discerned from our data directly or indirectly converge on the targets of Mef2. The combination of this evidence perhaps warrants greater attention to the spatio-temporal regulation of Mef2 during diapause, exploring its role as a central mediator of growth and aging in response to the environment.

### NELF-dependent RNAPII pausing in diapause

The prevalence of NELF and recruiters of NELF in our data implies a role for RNAPII pausing in adult diapause. RNAPII pausing is best characterized in *Drosophila* during early development as a preparatory mechanism driving the transcription of morphogenic ecdysone response genes (Mazina et al., 2021). However, recent work shows RNAPII pausing is more broadly detectable during later metamorphic checkpoints, as well (Mazina et al., 2022). This suggests RNAPII pausing is a general feature of tissue remodeling, particularly during phases like diapause wherein the organism anticipates later signals for further development. However, it has been shown that NELF is dispensable for RNAPII pausing. Aoi et al., 2020, showed that while NELF depletion does not enable transcript elongation from paused RNAPII, it induces advancement of the complex to a second pause point at the +1 nucleosome. This result indicates compound mechanisms to ensure effective pausing. NELF restricts transcription by interfering with RNAPII processivity: NELF binds the nascent RNA transcript, controls recruitment of the cap-binding and pre-initiation complexes, and competitively binds sites for positive elongation factors PAF (RNA Polymerase II-Associated Factor) and TFIIS (Transcription elongation factor IIS) (Yamaguchi et al., 2002; Aoi et al., 2020; Lahudkar et al., 2011; Vos et al., 2018). These functions of NELF constitute one mode of inhibition, though it appears that prevention of aberrant transcription may be achieved through multiple functionally redundant mechanisms, perhaps in a failsafe manner dependent on nucleosome distribution (Mazina et al., 2022). NELF is therefore dispensable, but an important “first pass” in coordinating multiple aspects of successful pausing. Relatedly, Myc has been demonstrated with the ability to release paused RNAPII in mouse embryonic stem cells. Assuming this function is conserved in *Drosophila*, the downregulation of Myc in our data may indicate one mode through which RNAPII pausing is preserved in the diapause state (Rahl et al., 2010).

## Materials and methods

### Survival assay

All flies used in this work were w^1118^ virgin females, reared at standard conditions (25°C, 12:12 light/dark, 40% relative humidity) on an agar-based diet supplemented with cornmeal (0.8%), sugar (10%), and yeast (2.5%). Newly eclosed virgin flies were identified by the presence of meconium. w^1118^ flystocks were obtained from Bloomington Stock Center. Flies were collected under mild CO2 and divided into plastic cages. Cages were then incubated at standard conditions or diapause-inducing conditions (11°C, 10:14 light/dark, 40% relative humidity) within eight hours post-eclosion. For flies kept at standard conditions, dead flies were counted and removed every 2-3 days coincident with the introduction of fresh food. For flies kept at diapause-inducing conditions, dead flies were counted and removed every 2-3 days, but food was replaced weekly due to slower food degradation and consumption rates at cooler temperatures. Flies were removed from diapause conditions on day 35 and placed at standard conditions to measure lifespan post-diapause. Visualization of data and statistical analysis between cohorts was conducted in R Statistical Software (v4.1.2; R Core Team 2022). The Kolmogorov-Smirnov test was used to determine whether survival data between cohorts had the same distribution.

### Diapause induction

Newly-eclosed virgin flies were collected under mild CO2 and divided into plastic cages. Cages were then placed into incubators housing four separate conditions within eight hours post-eclosion: standard conditions with 10 μL ethanol (99%) added every 2 days, standard conditions with 10 μL S-methoprene (39.4 mm, AK Scientific, effective dose 0.1 μg) added every 2 days, diapause-inducing conditions with 10 μL ethanol added every 2 days, and diapause-inducing conditions with 10 μL S-methoprene added every 2 days. S-methoprene was diluted in 99% ethanol. Dead flies were removed from cages at timepoints concurrent with ethanol or S-methoprene administration. Ethanol or S-methoprene was administered by adding 10 uL of the appropriate treatment to a cotton applicator. Applicators were suspended inside each cage, allowing the treatment to volatilize. Flies were fed from vials attached to the cages that contained the same food across conditions, including rearing conditions. Fresh vials were introduced weekly for flies at 11°C and every 2-3 days for flies at 25°C.

### ATAC-seq tissue and sequencing library preparation

Fat body tissue was dissected from 25-day-old females into minimal Schneider’s medium (Sigma-Aldrich). Prior to tissue collection, the diapause or non-diapause status was confirmed for each fly using observation of ovarian morphology. Only flies with the morphology expected for their respective conditions were dissected for fat body tissue; flies typically presented the expected morphology. Four biological replicates were collected per condition using fat body tissue from 25 flies per replicate. Nuclear extraction and library preparation protocols were adapted from Cources et al., 2017 and Davie et al., 2015. The NEBNext^™^ Illumina library quantification kit from New England Biosciences was used to measure final library concentration (Ref # E7630S, lot 10013445). The Nextera^™^ DNA library prep kit from Illumina (Ref # 15028212, lot 20256848) was used to tagment DNA, ligate adapter sequences and amplify fragments. The quality of ATAC-seq libraries was assessed via Bioanalyzer (Agilent) prior to sequencing.

### RNA-seq tissue and sequencing library preparation

Fat body tissue was dissected from 25-day-old females into Trizol (Ambion). Ovarian morphology was observed to confirm diapause status as described for our ATAC-seq tissue collection. Six replicates were collected per condition using fat body tissue from 25 flies per replicate. Tissue was then homogenized with steel beads and total RNA was extracted using the Direct-zol^™^ RNA MicroPrep kit from Zymo Research. RNA concentration and purity was assessed via Nanodrop^™^ and Qubit^™^ technologies prior to library preparation, which was completed at Novogene facilities. Briefly, mRNA was isolated from total RNA via using magnetic beads coated with polyT oligomers. mRNA was then fragmented and cDNA synthesized, followed by end repair, A-tailing, Illumina adapter ligation, size selection, PCR amplification and purification. The final concentrations and purity of libraries were then assessed via Qubit^™^ and qPCR. Fragment size distribution was assessed via bioanalyzer.

### Initial processing of ATAC-seq data

ATAC-seq libraries were pooled and sequenced in multiplex with a paired-end read length of 150 bp at an average depth of approximately 26 million reads per sample. The quality of raw reads was assessed using FastQC (http://www.bioinformatics.babraham.ac.uk/projects/fastqc). Adapters were trimmed using Trimgalore! (https://github.com/FelixKrueger/TrimGalore) and Cutadapt (Martin, 2011) from the Bioinformatics Group at the Babraham Institute. Trimmed reads were then aligned to the Flybase dm6 reference genome using Bowtie2 with the parameter --no-discordant and a maximum fragment length of 1000 bp (dos Santos et al., 2015; Langmead & Salzberg, 2012). Mitochondrial reads were removed from ATAC-seq files using base Unix and Samtools (Danecek et al., 2021). Resulting sam files were converted to bam via Samtools. Picard (http://broadinstitute.github.io/picard/)and Samtools were then used to remove optical duplicate reads and then index the filtered bam files. Peaks were then called from filtered bam files using MACS version 2 with a minimum FDR cutoff of 0.05 and parameters --nomodel, --shift −100, --extsize 200, --keep-dup-all and–call-summits (Zhang et al., 2008). Blacklisted regions were removed from ATAC-seq files using Bedtools intersect (Quinlan & Hall, 2010).

### Initial processing of RNA-seq data

RNA-seq libraries were pooled and sequenced in multiplex with a paired-end read length of 150 bp at an average depth of approximately 52 million reads per sample. The quality of raw reads was assessed using FastQC and quality reports from Novogene. Alignment and initial processing for RNA-seq data proceeded using similar steps as described here for ATAC-seq, but peaks were not called and mitochondrial reads were retained. Bam files were then exported for further analysis in R.

### Differential accessibility analysis

ATAC-seq output from MACS2 was analyzed for differential accessibility between conditions using DiffBind in R (Stark & Brown, 2011; Ross-Ines et al., 2012). To arrive at the subset of consensus regions shared between both 11°C conditions while excluding those regions that are also accessible at 25°C, the dba.contrast function was first used to compare both 11°C conditions. The dba.report function was then used to derive the set of regions that are not differentially accessible between either 11°C treatment via the argument bNotDB = TRUE and a p value cutoff of 0.01 (set 1). A consensus set from all 25°C samples was then created using the dba.peakset function, specifying a minimum overlap of 0 bp to include regions that may differ significantly by treatment (set 2). Set 2 was then intersected with set 1 to arrive at the set of regions shared among all four conditions, as well as the set of regions that were exclusive to both 11°C conditions (accessible in diapause) and those that were exclusive to either 25°C conditions (inaccessible in diapause). Differential accessibility was defined with an FDR cutoff of 0.01.

### Differential expression analysis

Gene counts for RNA-seq were obtained from processed bam files using the Flybase dm6 transcript set and featureCounts, a program within the RSubread package (Liao et al., 2014; Liao et al., 2019). Multi-mapping reads were not counted. Counts were then differentially analyzed by temperature and treatment using DESeq2 and the design formula ~ treatment + temperature. Data normalization was performed using the rlog() function in base R. Differentially expressed genes with an FDR-adjusted p value of less than 0.01 were considered significantly differentially expressed.

### Genomic enrichment analysis

Genomic feature enrichment of differentially accessible loci was calculated using the R package ChIPseeker while referencing the dm6 annotation packages from UCSC and Carlson, 2019. Example loci were visualized using Gviz (Hahne & Ivanek, 2016).

### Motif enrichment analysis

Motif enrichment analysis was performed using i-cisTarget under default parameters: a minimum normalized enrichment score of 3 and a minimum overlap of 40% with candidate database motifs were required to be considered a match with input regions. Motif enrichment analysis was additionally performed using HOMER to compare with results from i-cisTarget using default parameters and a fragment size of 200 bp.

### STRING interaction network analysis

Transcription factor nodes and edge counts were derived using the STRINGdb R package, database version 11.5 (Szklarczyk et al., 2021). Network plots were generated and merged in Cytoscape version 3.9.1 (Shannon et al., 2003) using the plugins enhancedGraphics (1.5.4), clusterMaker2 (2.2), cytoHubba (0.1), stringApp (1.7.1), Pathway Scoring Application (3.0.2), yFiles Layout Algorithms (1.1.2), and Largest Subnetwork (1.3). Transcription factors were derived from interaction data by referencing lists from Flybase (release FB2022_03) and Flymine (version 53). GAF interaction data was added from Lomaev et. al, 2017.

### Pathway and overrepresentation analysis

SPIA was performed using the Graphite web implementation under default parameters using the Reactome pathway database (Draghici et al., 2007; Tarca et al., 2009; Sales et al., 2013; Gillespie et al., 2022). Gene Ontology (GO) analysis was performed using the clusterProfiler package in R with a q value cutoff of 0.05 and the Benjamini-Hochberg p value adjustment procedure (Ashburner et al., 2000; Wu et al., 2021; Gene Ontology Consortium, 2021). Revigo was used to consolidate GO terms from clusterProfiler with a similarity cutoff value of 0.5. P values were associated with GO terms when consolidating data in Revigo. Co-regulation analysis was performed using the R implementation of FlyEnrichr under default parameters and the Drosophila Interactions Database (DroID, version 2015) (Chen et al., 2013; Kuleshov et al., 2016). Pathway analysis was also performed using FlyEnrichR under default parameters with the Kyoto Encyclopedia of Genes and Genomes (KEGG, version 2019). GSEA (Subramanian et al., 2005; Mootha et al., 2003) was performed using gene matrices from KEGG (Kanehisa & Goto, 2000; Kanehisa, 2019; Kanehisa et al., 2021) and Reactome, which were generated for *Drosophila* by Cheng et al., 2021.

### Immunocytochemistry and EdU incorporation

Protocol for immunostaining of adult fat body tissue was adapted from Gouge and Christensen, 2010, as well as the manual for Thermo Fisher’s Click-iT^™^ EdU Cell Proliferation Kit for Imaging (Ref # C10337). Briefly, dissected fat body tissue was suspended in minimal Schneider’s medium with 30 μM EdU for 30 minutes at room temperature (RT). Tissue was fixed in 3.7% formaldehyde and permeabilized with 0.1% Triton X-100. Tissue was then blocked in 10% normal goat serum for 10 minutes and subsequently incubated in anti-Delta serum (C594.9B, Developmental Studies Hybridoma Bank) for 2 hours at RT. After washing in PBS, tissue was incubated in Cy5-conjugated secondary antibody for another 2 hours at RT. Tissue was washed and suspended in Click-iT reaction cocktail, then incubated 30 minutes in darkness. Hoechst was used to label nuclei. Tissue was mounted in 80% glycerol and imaged using the Olympus VS200 Slide Scanner. Imaged were analyzed in Olympus software OlyVIA (version 3.4.1) and ImageJ (version 1.53s). This protocol is available as a supplement (supplemental protocol 1.)

### Publicly available ChIP-seq data

ChIP-seq data from juvenile flies were acquired from modENCODE (Celniker et al., 2009) under identification numbers 5102 and 5097. Raw sequencing files were aligned to the Flybase dm6 reference genome and further processed using the method described here for ATAC-seq data. Background peaks were removed within the peak calling function of MACS2. Background-less files were then output as bedGraph and converted to bigWig format using the UCSC function bedGraphToBigWig. Files were then compared to differentially accessible promoter regions derived from ATAC-seq via ChIPseeker. The deepTools computeMatrix function was used under reference point mode, a window size of ±5 kb, and the skipZeros parameter. deepTools plotHeatmap was used with default parameters (Ramírez et al., 2014).

### Accession numbers

Raw sequencing files and processed bam files are available for ATAC-seq and RNA-seq via NCBI-GEO (http://www.ncbi.nlm.nih.gov/geo/) under accession numbers GSE212862 and GSE212863.

## Supporting information

List of supplemental tables

Supplemental protocol 1

## Acknowledgements

We thank Jacqueline Howells for insightful discussions around this work. We additionally acknowledge the guidance of Jackson Taylor, Erica Larschan, John Sedivy, Nicola Neretti, Bérénice Benayoun and Marc Tatar. This work was supported by funding from 2T32AG041688-06 (C.H.) and 2R37AG024360-11 (NIH NIA).

