## Supplemental figures for "Multiomic Analysis of Adult Diapause in *Drosophila melanogaster* Identifies Hallmarks of Cellular Quiescence"

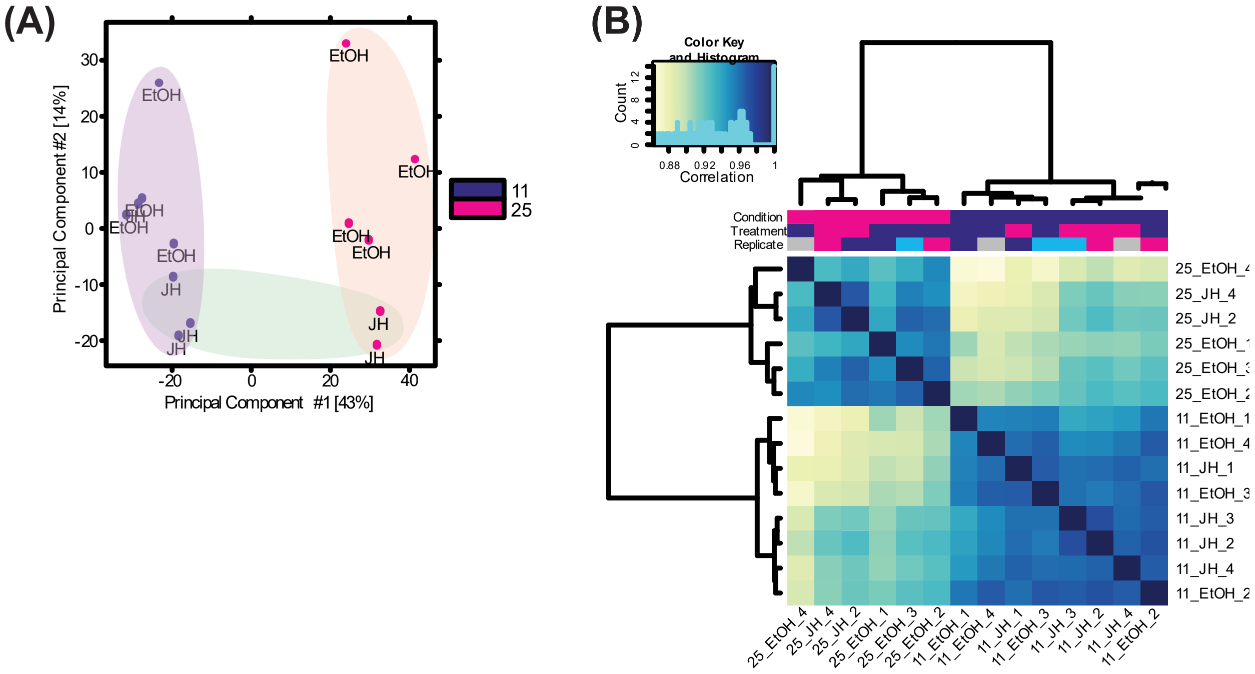


Figure S.1: Correlation between ATAC-seq replicates

**(A)** Principal component analysis of ATAC-seq replicates. Samples clustered primarily by temperature. **(B)** Pearson correlation heatmap comparing all ATAC-seq replicates. Samples again clustered primarily by temperature.

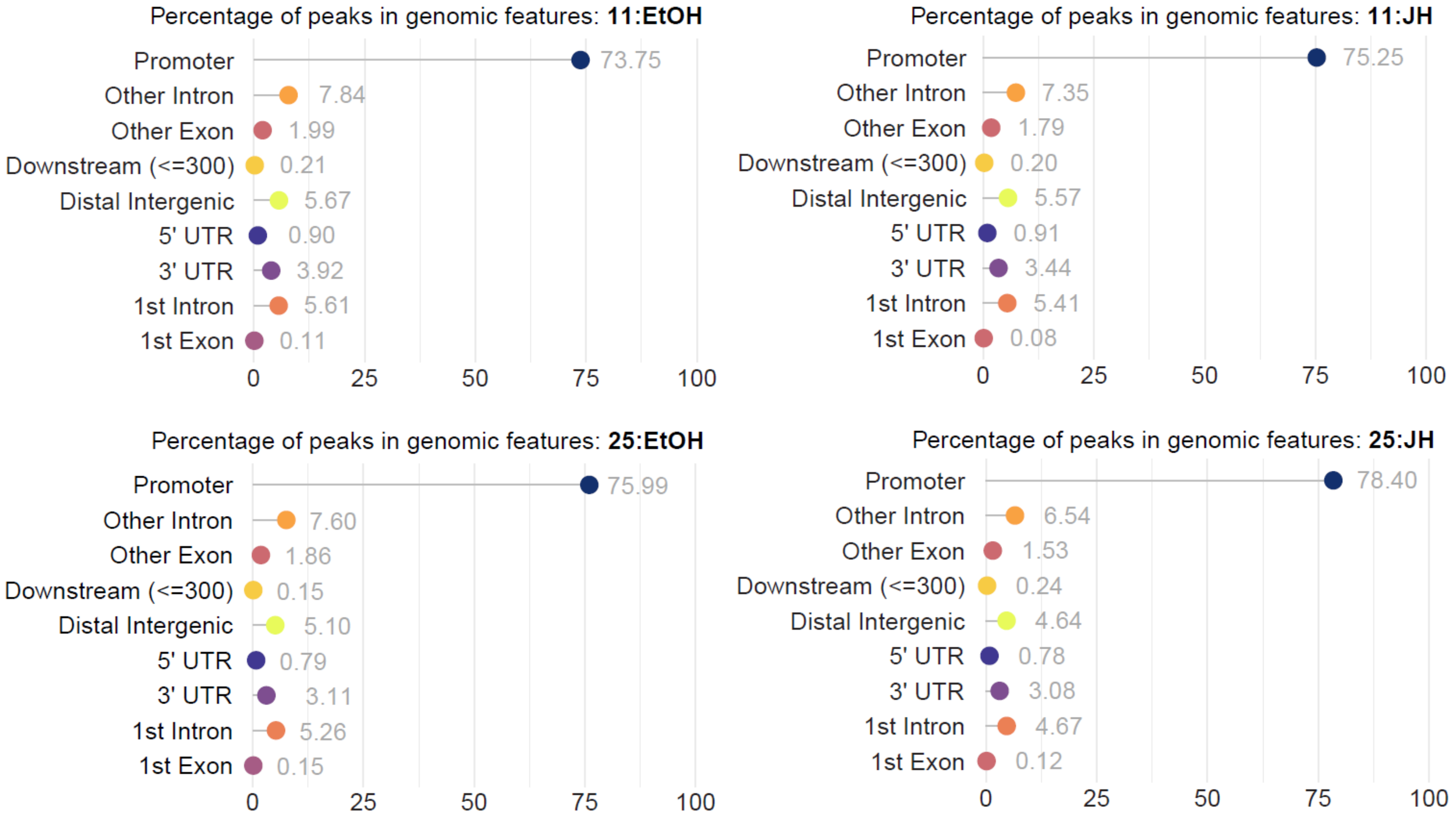


Figure S.2: Euchromatin is distributed similarly within genomic features across conditions

**(A)** Distribution of accessible loci per condition, before differential analysis, according to the genomic features they map to. Promoters were defined within ±1 kb of a transcription start site.

### Homer Known Motif Enrichment Results (accessible, FDR<0.01)

Total Target Sequences = 513, Total Background Sequences = 45095

| Rank | Motif | Name | P-value | log P-pvalue | q-value (Benjamini) | % of Targets Sequences with Motif | % of Background Sequences with Motif |
|---|---|---|---|---|---|---|---|
| 1 | 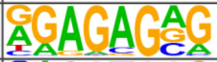 | Trl(Zf)/S2-GAGAfactor-ChIP-Seq(GSE40646)/Homer | 1e-22 | -5.113e+01 | 0.0000 | 36.26% | 17.94% |
| 2 | 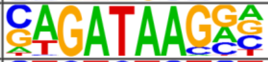 | Gata1(Zf)/K562-GATA1-ChIP-Seq(GSE18829)/Homer | 1e-11 | -2.672e+01 | 0.0000 | 14.23% | 5.81% |
| 3 | 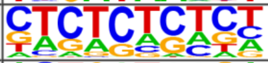 | GAGA-repeat/Arabidopsis-Promoters/Homer | 1e-11 | -2.664e+01 | 0.0000 | 15.59% | 6.72% |
| 4 | 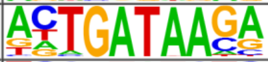 | PQM-1(?)/cElegans-L3-ChIP-Seq(modEncode)/Homer | 1e-10 | -2.379e+01 | 0.0000 | 11.11% | 4.20% |
| 5 | 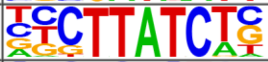 | Gata2(Zf)/K562-GATA2-ChIP-Seq(GSE18829)/Homer | 1e-9 | -2.226e+01 | 0.0000 | 14.04% | 6.28% |
| 6 | 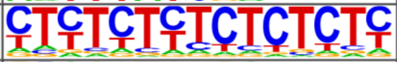 | GAGA-repeat/SacCer-Promoters/Homer | 1e-9 | -2.127e+01 | 0.0000 | 43.47% | 30.61% |
| 7 | 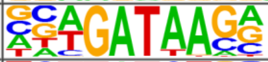 | Gata4(Zf)/Heart-Gata4-ChIP-Seq(GSE35151)/Homer | 1e-8 | -2.024e+01 | 0.0000 | 19.69% | 10.72% |
| 8 | 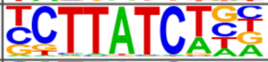 | Gata6(Zf)/HUG1N-GATA6-ChIP-Seq(GSE51936)/Homer | 1e-8 | -1.955e+01 | 0.0000 | 17.54% | 9.24% |
| 9 | 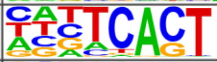 | STZ(C2H2)/colamp-STZ-DAP-Seq(GSE60143)/Homer | 1e-7 | -1.819e+01 | 0.0000 | 63.55% | 51.24% |
| 10 | 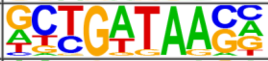 | Unknown5/Drosophila-Promoters/Homer | 1e-6 | -1.611e+01 | 0.0000 | 14.42% | 7.60% |
| 11 | 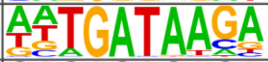 | ELT-3(Gata)/cElegans-L1-ELT3-ChIP-Seq(modEncode)/Homer | 1e-6 | -1.550e+01 | 0.0000 | 9.75% | 4.39% |
| 12 | 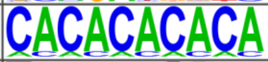 | SeqBias: CA-repeat | 1e-6 | -1.467e+01 | 0.0000 | 64.33% | 53.48% |
| 13 | 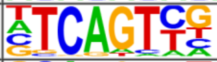 | Initiator/Drosophila-Promoters/Homer | 1e-5 | -1.261e+01 | 0.0003 | 28.27% | 19.89% |
| 14 | 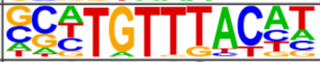 | FOXK2(Forkhead)/U2OS-FOXK2-ChIP-Seq(E-MTAB-2204)/Homer | 1e-5 | -1.259e+01 | 0.0003 | 12.28% | 6.72% |
| 15 | 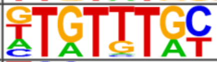 | PHA-4(Forkhead)/cElegans-Embryos-PHA4-ChIP-Seq(modEncode)/Homer | 1e-5 | -1.162e+01 | 0.0006 | 53.80% | 44.26% |
| 16 | 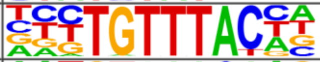 | FOXP1(Forkhead)/H9-FOXP1-ChIP-Seq(GSE31006)/Homer | 1e-4 | -1.147e+01 | 0.0006 | 10.14% | 5.36% |
| 17 | 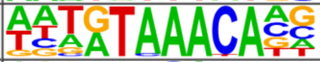 | FoxL2(Forkhead)/Ovary-FoxL2-ChIP-Seq(GSE60858)/Homer | 1e-4 | -1.115e+01 | 0.0008 | 18.71% | 12.20% |
| 18 | 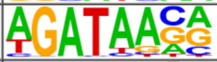 | GATA3(Zf)/iTreg-Gata3-ChIP-Seq(GSE20898)/Homer | 1e-4 | -1.101e+01 | 0.0009 | 22.22% | 15.21% |
| 19 | 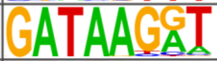 | At3g11280(MYBrelated)/col-At3g11280-DAP-Seq(GSE60143)/Homer | 1e-4 | -1.070e+01 | 0.0012 | 16.18% | 10.24% |
| 20 | 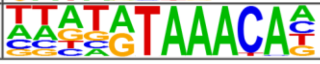 | Foxf1(Forkhead)/Lung-Foxf1-ChIP-Seq(GSE77951)/Homer | 1e-4 | -1.060e+01 | 0.0012 | 20.47% | 13.85% |

Figure S.3: Top 20 motif enrichment results for genomic loci that become accessible in diapause (HOMER)

HOMER motif enrichment analysis was performed on genomic loci that become accessible in diapause using default parameters and a fragment size of 200. HOMER returned results with less species specificity than i-cisTarget, though certain highly significant results are the same between platforms.

### Homer Known Motif Enrichment Results (inaccessible, FDR<0.01)

Total Target Sequences = 692, Total Background Sequences = 46923

| Rank | Motif | Name | P-value | log P-pvalue | q-value (Benjamini) | % of Targets Sequences with Motif | % of Background Sequences with Motif |
|---|---|---|---|---|---|---|---|
| 1 | 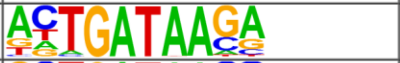 | PQM-1(?)/cElegans-L3-ChIP-Seq(modEncode)/Homer | 1e-35 | -8.138e+01 | 0.0000 | 18.06% | 4.91% |
| 2 | 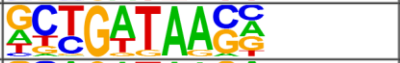 | Unknown5/Drosophila-Promoters/Homer | 1e-28 | -6.670e+01 | 0.0000 | 21.97% | 8.07% |
| 3 | 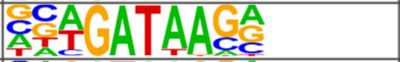 | Gata4(Zf)/Heart-Gata4-ChIP-Seq(GSE35151)/Homer | 1e-26 | -6.165e+01 | 0.0000 | 27.60% | 12.26% |
| 4 | 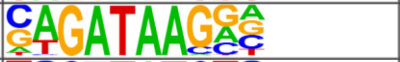 | Gata1(Zf)/K562-GATA1-ChIP-Seq(GSE18829)/Homer | 1e-23 | -5.459e+01 | 0.0000 | 17.77% | 6.41% |
| 5 | 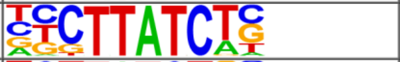 | Gata2(Zf)/K562-GATA2-ChIP-Seq(GSE18829)/Homer | 1e-22 | -5.219e+01 | 0.0000 | 18.35% | 6.96% |
| 6 | 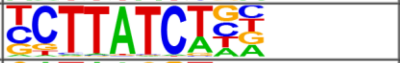 | Gata6(Zf)/HUG1N-GATA6-ChIP-Seq(GSE51936)/Homer | 1e-22 | -5.200e+01 | 0.0000 | 23.70% | 10.51% |
| 7 | 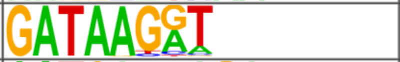 | At3g11280(MYBrelated)/col-At3g11280-DAP-Seq(GSE60143)/Homer | 1e-17 | -4.060e+01 | 0.0000 | 22.83% | 11.17% |
| 8 | 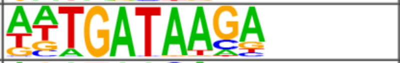 | ELT-3(Gata)/cElegans-L1-ELT3-ChIP-Seq(modEncode)/Homer | 1e-17 | -3.968e+01 | 0.0000 | 14.60% | 5.65% |
| 9 | 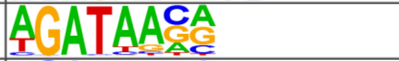 | GATA3(Zf)/iTreg-Gata3-ChIP-Seq(GSE20898)/Homer | 1e-16 | -3.730e+01 | 0.0000 | 30.35% | 17.42% |
| 10 | 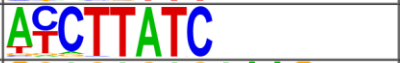 | At5g05790(MYBrelated)/col-At5g05790-DAP-Seq(GSE60143)/Homer | 1e-15 | -3.589e+01 | 0.0000 | 23.12% | 11.99% |
| 11 | 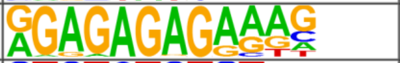 | FRS9(ND)/col-FRS9-DAP-Seq(GSE60143)/Homer | 1e-12 | -2.794e+01 | 0.0000 | 5.92% | 1.55% |
| 12 | 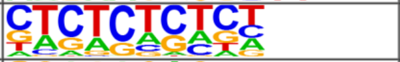 | GAGA-repeat/Arabidopsis-Promoters/Homer | 1e-11 | -2.646e+01 | 0.0000 | 13.58% | 6.29% |
| 13 | 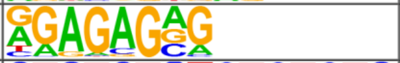 | Trl(Zf)/S2-GAGAfactor-ChIP-Seq(GSE40646)/Homer | 1e-8 | -2.070e+01 | 0.0000 | 26.01% | 16.90% |
| 14 | 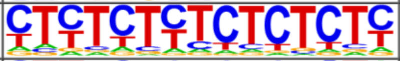 | GAGA-repeat/SacCer-Promoters/Homer | 1e-7 | -1.786e+01 | 0.0000 | 45.09% | 34.85% |
| 15 | 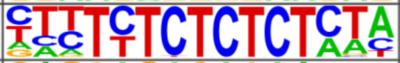 | BPC6(BBRBPC)/col-BPC6-DAP-Seq(GSE60143)/Homer | 1e-6 | -1.584e+01 | 0.0000 | 3.03% | 0.75% |
| 16 | 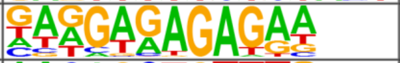 | BPC1(BBRBPC)/colamp-BPC1-DAP-Seq(GSE60143)/Homer | 1e-5 | -1.359e+01 | 0.0001 | 6.65% | 3.06% |
| 17 | 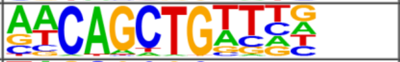 | E-box/Drosophila-Promoters/Homer | 1e-5 | -1.225e+01 | 0.0003 | 4.77% | 1.97% |
| 18 | 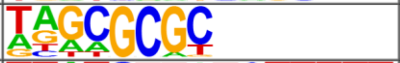 | DPL-1(E2F)/cElegans-Adult-ChIP-Seq(modEncode)/Homer | 1e-4 | -1.005e+01 | 0.0023 | 16.91% | 11.78% |
| 19 | 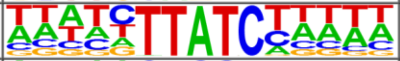 | At5g58900(MYBrelated)/colamp-At5g58900-DAP-Seq(GSE60143)/Homer | 1e-3 | -8.359e+00 | 0.0120 | 23.12% | 17.80% |
| 20 | 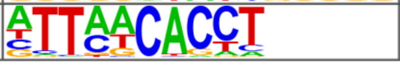 | Eomes(T-box)/H9-Eomes-ChIP-Seq(GSE26097)/Homer | 1e-3 | -7.806e+00 | 0.0199 | 25.29% | 19.99% |

Figure S.4: Top 20 motif enrichment results for genomic loci that become inaccessible in diapause (HOMER)

HOMER motif enrichment analysis was performed on genomic loci that become inaccessible in diapause using default parameters and a fragment size of 200. HOMER returned results with less species specificity than i-cisTarget, though certain highly significant results are the same between platforms.

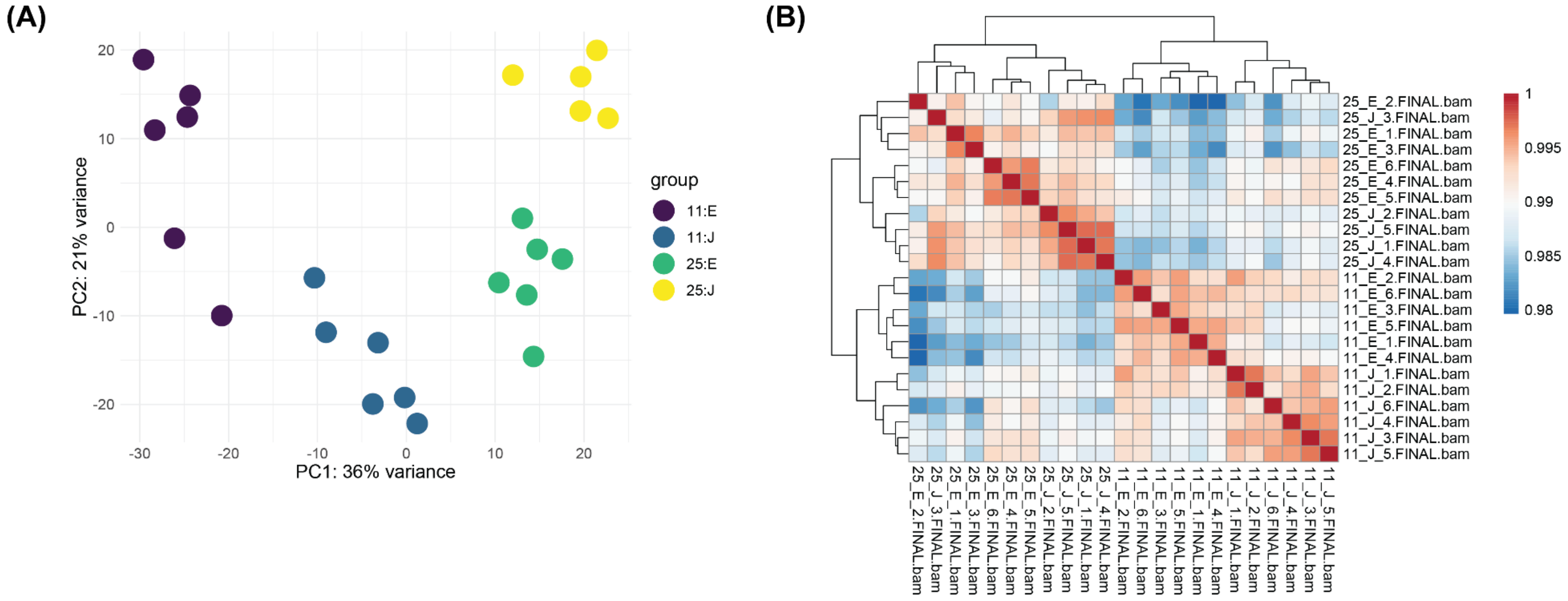


Figure S.5: Correlation between RNA-seq replicates

**(A)** Principal component analysis of RNA-seq replicates. Samples clustered visibly by condition. **(B)** Pearson correlation heatmap comparing all RNA-seq replicates. Samples clustered by condition.

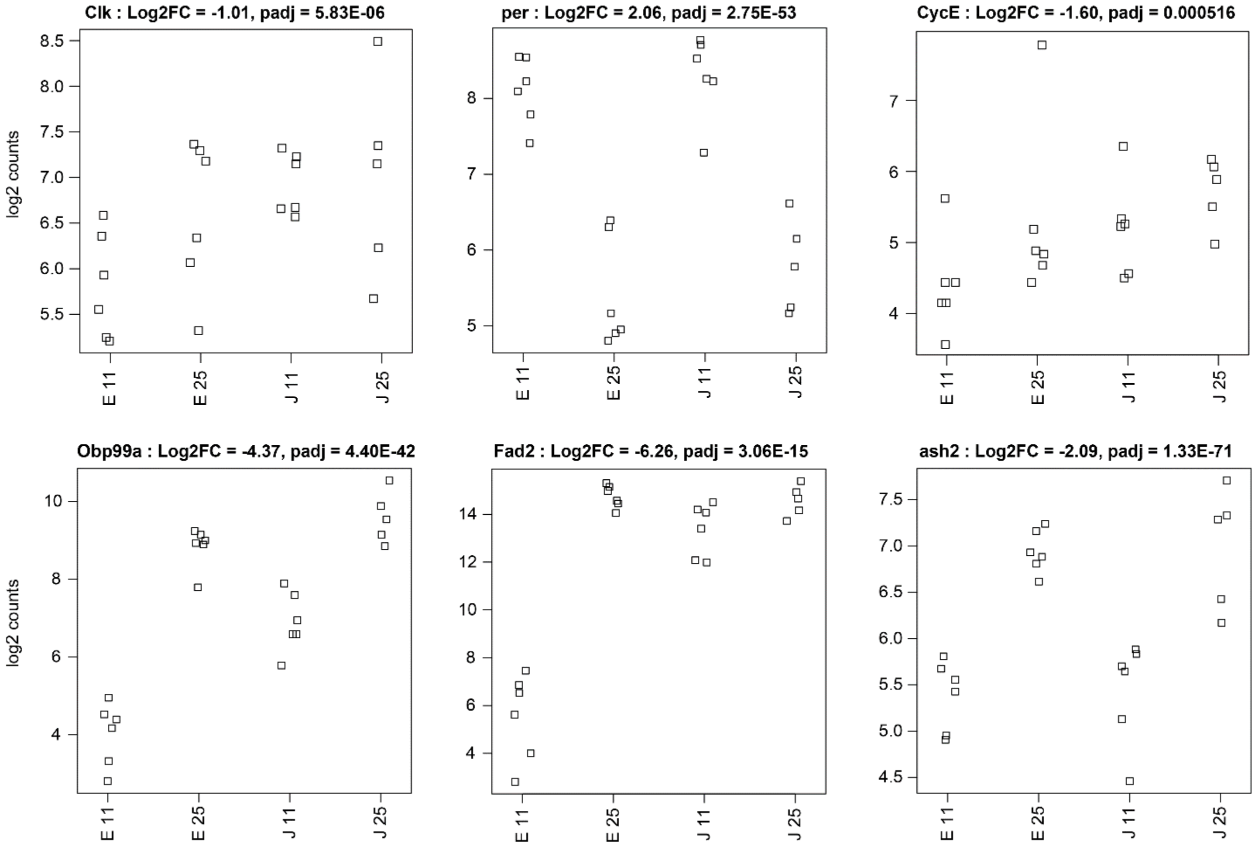


Figure S.6: Highlighted differentially expressed genes

Divergent transcriptional regulation of *clk* and *per* are demonstrative results for the successful induction of diapause. *clk* transcription appears more dependent on JH status than *per. Obp99a* and *Fad2* are known to be upregulated in response to JH and are here presented as demonstrative results for the success of our JH supplementation experimental design. Ash2 is a catalytic component of the Trithorax nucleosome remodeling complex. *cycE* appears to be downregulated in a JH-dependent manner.

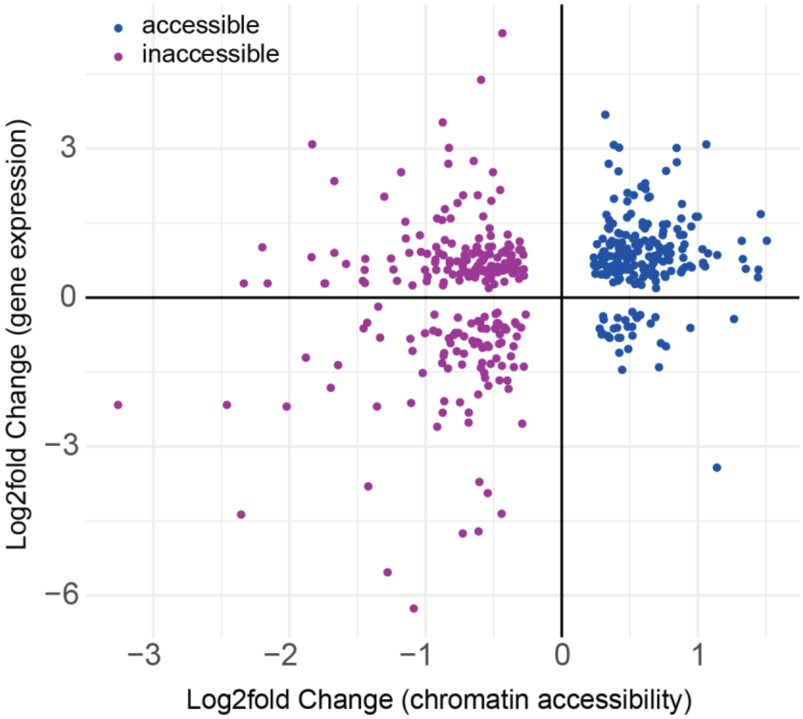


Figure S.7: Comparison of magnitudes in gene expression and chromatin accessibility

A comparison between the log2fold changes in gene expression and the log2fold change in chromatin accessibility for those DEGs with differentially accessible chromatin. Correlation between magnitude of chromatin opening/closing and gene up/downregulation would present as a datapoints fitting an x=y distribution. However, we observed no trend in magnitudes, indicating the magnitude of a change in chromatin accessibility has no effect on the level of gene expression observed at that locus.
