## Supplementary material for "Multiomic Analysis of Adult Diapause in *Drosophila melanogaster* Identifies Hallmarks of Cellular Quiescence": List of supplemental tables

Below are downloadable links to supplemental tables.

*Table S1.* *ATAC-seq pearson correlations per replicate per condition*
*Table S2.* *Differentially accessible loci*
*Table S3.* *Results from HOMER motif analysis of accessible loci in diapause*
*Table S4.* *Results from HOMER motif analysis of inaccessible loci in diapause*
*Table S5.* *Results from i-cisTarget motif analysis of accessible loci in diapause*
*Table S6.* *Results from i-cisTarget motif analysis of inaccessible loci in diapause*
*Table S7.* *GO terms derived from genes with accessible promoters*
*Table S8.* *GO terms derived from genes with inaccessible promoters*
*Table S9.* *Upstream regulatory factors derived from genes with accessible promoters*
*Table S10.* *Upstream regulatory factors derived from genes with inaccessible promoters*
*Table S11.* *Interaction data for all genes with accessible promoters*
*Table S12.* *Interaction data for all genes with inaccessible promoters*
*Table S13.* *Differentially expressed genes*
*Table S14.* *Results from SPIA performed on differentially expressed genes, p.adjust<0.01*
*Table S15.* *Results from SPIA performed on differentially expressed genes, p.adjust<0.05*
*Table S16.* *Interaction data for differentially expressed transcription factors*
*Table S17.* *Upstream regulatory factors derived from upregulated genes*
*Table S18.* *Upstream regulatory factors derived from downregulated genes*
*Table S19.* *GO terms derived from differentially expressed genes with differentially accessible chromatin*
*Table S20.* *Results from SPIA performed on differentially expressed genes with differentially accessible chromatin, p.adjust<0.05*
