## Supplemental protocol 1 for "Multiomic Analysis of Adult Diapause in *Drosophila melanogaster* Identifies Hallmarks of Cellular Quiescence"

**EdU labeling of dissected adult fat body (*Drosophila melanogaster*)**

*Preparation*

30uM solution of EdU in Schneider’s medium, allow to reach room temp

- 1.5 uL of 10mM EdU in 498.5 uL Schneider’s medium, scale as needed

9-well plate with 50 uL Schneider’s medium per well

10 mL 3% BSA in 1x PBS (1.5 g in 50 mL, or 0.3 g in 10 mL)

3.7% formaldehyde in 1x PBS (50 uL/well)

0.1% Triton-X-100 in 1x PBS (50 uL/well)

*During the Triton-X-100 incubation step:* [Click-iT reaction cocktail](https://www.thermofisher.com/document-connect/document-connect.html?url=https%3A%2F%2Fassets.thermofisher.com%2FTFS-Assets%2FLSG%2Fmanuals%2Fmp10338.pdf) (100 uL/well):

Assuming 500 uL final volume:

(1x Click-iT Reaction Buffer: 387 uL distilled water + 43 uL stock buffer)

- 1× Click-It Reaction Buffer	430 uL
- CuSO4	20 uL

(Reaction Buffer Additive: 45 uL distilled water + 5 uL buffer additive)

- Alexa Fluor Azide (diluted in DMSO per kit instruction)	1 uL
- Reaction Buffer Additive	50 uL

*Add reagents in THIS ORDER

*Use this cocktail within 15 minutes of preparation

5 ug/mL Hoescht33342 in 1x PBS (50 uL/well)

1. Make 3% BSA, prepare EdU solution, set items to thaw (primary Ab, Schneider’s, aphidicolin, NGS, EdU), prepare dissection platform & distribute medium into wells
2. Dissect tissue into medium
3. Treat negative control samples with 100 ug/mL aphidicolin for 15 minutes
   - At stock = 1 mg/mL, add 5 uL of stock to every 50 uL of medium
4. Remove solution from wells, wash twice with 100 uL 3% BSA
5. Remove BSA, add 50 uL Schneider’s medium and 50 uL EdU solution, incubate 30 minutes at room temp
6. Remove EdU solution, wash twice with 100 uL of 3% BSA
7. Remove BSA, add 50 uL of 3.7% formaldehyde, incubate 5 minutes
8. Remove formaldehyde, wash twice in 3% BSA
9. Remove BSA, add 100 uL of 0.1% Triton-X-100, incubate 20 minutes (dilute Abs now)
10. Remove Triton-X-100, wash twice in 3% BSA
11. Block in 10% NGS in PBS for 10 minutes
12. Remove NGS, wash 3x in PBST
13. Add 50 uL of primary Ab solution (diluted in PBST), incubate 2 hours RT
14. Remove antibody (store with sodium azide 1:1000), wash 3x in PBS
15. Add 50 uL secondary Ab (diluted in PBST), incubate 2 hours RT
    - Prepare Click-iT reaction cocktail near the end of this incubation period
16. Remove antibody, wash 3x in BSA
17. Remove BSA, add 100 uL of Click-iT reaction cocktail, incubate 30 minutes in darkness

- Keep samples in darkness for the remainder of the protocol

18. Remove Click-iT reaction cocktail, wash twice in 3% BSA
19. Remove BSA, add 50 uL Hoescht solution, incubate 15 minutes
20. Remove Hoescht solution, wash twice with 1x PBS
21. Add 5 uL of 1x PBS to slides, place tissue on slides. Remove excess PBS after transfer.
22. Mount tissue on slides with 10 uL 80% glycerol, place coverslip
23. Visualize using fluorescence or confocal microscope
